# Strong gene flow across an urbanised coastal landscape in a dune specialist digger wasp

**DOI:** 10.1101/2023.04.15.537020

**Authors:** Femke Batsleer, Fabien Duez, Dirk Maes, Dries Bonte

**Affiliations:** Ghent University, Department of Biology, Terrestrial Ecology Unit, Karel Lodewijk Ledeganckstraat 35, 9000 Gent, Belgium; Institut des Sciences de l’Evolution, Université de Montpellier, CNRS, IRD, Montpellier, France; Research Institute for Nature and Forest (INBO), Herman Teirlinckgebouw, Havenlaan 88 bus 73, 1000 Brussel, Belgium; Radboud Institute for Biological and Environmental Sciences (RIBES), Radboud University, PO Box 9010, NL-6500 GL Nijmegen, The Netherlands

**Keywords:** landscape genetics, landscape resistance modelling, resistanceGA, resistance optimisation, dispersal, Hymenoptera

## Abstract

**Context:** Genetic connectivity is often disrupted by anthropogenic habitat fragmentation, and therefore often a focus in landscape-scale conservation. Landscape genetics methods allow for studying functional connectivity in heterogenous landscapes in detail and have the potential to inform conservation measures for a species’ regional persistence. Yet, for insects, functional connectivity through landscape genetics remains largely unexplored.

**Objectives:** We aimed to assess the functional connectivity in the dune-specialist digger wasp *Bembix rostrata*, in a human-altered 15 km section along the Belgian coast, based on landscape genetics methods.

**Methods:** We optimised landscape resistance distances according to individual genetic distances with the package ResistanceGA. We combined this with a multi-model inference approach to deduce relative conductance or resistance of gene flow to natural and anthropogenic landscape types.

**Results:** Overall, the populations of this dune-specialist insect are genetically well-connected. We detected— on top of the prominent background process of isolation-by-distance—a weak but consistent signal of urban features facilitating gene flow.

**Conclusions:** Urban areas are not a barrier to genetic connectivity of the dune-specialist insect *B. rostrata* in the focal human-altered landscape. However, because urbanisation leads to larger scale fragmentation, its impact on the distribution of populations in the landscape and related effective regional gene flow remains substantial. As this species depends on early-succession dune vegetations, restoring and increasing sand dynamics at the local and landscape scale should be the focus of conservation aimed at the regional species’ persistence, rather than trying to increase habitat connectivity in this human-altered dune ecosystem.

## Introduction

Movement of individuals through landscapes is an inherent component of dispersal and can lead to exchange between populations, resulting in gene flow and connectivity. Connectivity can ultimately support a species’ regional persistence, as it maintains or supports a wide range of natural processes, such as resource use, demographic rescue, genetic diversity, range expansion, or metapopulation dynamics (Hanski 1998; Crooks and Sanjayan 2006; McRae et al. 2008). The environment in between habitat patches (i.e. the matrix) can impede or facilitate the movement or dispersal of an organism. This matrix is in reality more heterogenous than the practical simplification of a homogeneous matrix, especially in human-altered landscapes (Hein et al. 2003; Manel and Holderegger 2013). The realised connectivity of a species is influenced by distances between populations and the specific landscape context: the structural connectivity. However, it also depends heavily on the behavioural responses of an organism to different components of the landscape (Hodgson et al. 2022). These responses vary inherently between species and individuals (Baguette and Van Dyck 2007; Knowlton and Graham 2010; Logghe et al. 2024). The integration of movement into measures of functional connectivity (Tischendorf & Fahrig, 2003) is, therefore, crucial for understanding the dynamics of spatially structured populations (or metapopulations) and to inform effective conservation measures (McRae 2006).

A general view on functional connectivity is that many organisms will avoid unfamiliar habitat altogether—they exhibit high reluctance to cross gaps and favour instead considerable detours— making unfamiliar habitat a barrier to movement (Develey and Stouffer 2008; Knowlton and Graham 2010; Driscoll et al. 2013). This is the general assumption in for instance least cost mapping, which considers only the most cost-effective path through a landscape (Adriaensen et al. 2003). Unfamiliar habitat can, however, also act as a facilitator to movements when search behaviour of individuals in unfamiliar habitat is more directional—and thus faster—than in favoured habitat (Van Dyck and Baguette 2005; Knowlton and Graham 2010). Or conversely, movements in more suitable habitats can be impeded because of the availability of resources, leading to less directed—and thus slower— exploratory, routine movements than in unfamiliar habitat (Van Dyck and Baguette 2005; McRae et al. 2008; Keller et al. 2012). The behavioural differences between more directional movement paths (covering greater distances) during displacement and (slower) exploratory, more random, routine movements are for insects best studied in butterfly species (Schtickzelle et al. 2006; Kuefler et al. 2010). Independent of the exact mechanism behind this potential (relative) increase in movement through the matrix, the eventual functional connectivity will remain dependent on the distance between habitats and thus the scale of the fragmentation. The same matrix typology or landscape category can therefore potentially facilitate or constrain connectivity in different landscapes, depending on the eventual distance-related costs in terms of survival and energy-use (Segelbacher et al. 2010; Spear et al. 2010; Bonte et al. 2012; Richardson et al. 2016; Haran et al. 2017).

How a landscape influences the mobility of an organism has implications for the design of regional conservation measures. If animals show high reluctance towards crossing habitat boundaries, creating corridors might be an effective structural connectivity measure to increase functional connectivity between subpopulations. Such regional landscape-scale conservation measures, including corridors and stepping-stones, which are based on logical and sound scientific principles, have been widely embraced by conservation practitioners and policy-makers (Crooks and Sanjayan 2006; Watts et al. 2016). However, there is much debate about the effectiveness of landscape-scale conservation and the emphasis on increasing connectivity for species’ regional persistence in conservation decisions (Hodgson et al. 2009; Humphrey et al. 2015; Watts et al. 2016). The emphasis of practitioners and policy-makers on landscape-scale conservation is probably biased by a focus towards larger terrestrial vertebrates, for which structural connectivity—such as corridors—seems effective (Humphrey et al. 2015). Empirical evidence for many other groups is more limited and equivocal (Ovaskainen et al. 2008; Watts et al. 2016). In insects, it is generally assumed that the regional persistence of spatially structured populations is mostly determined by local demography (Richardson et al. 2016; Drake et al. 2022).

In such cases, site-based conservation focusing on improving habitat area and quality could be more effective than increasing structural connectivity on the landscape-scale. To effectively direct the limited resources available for conservation measures towards implementations with the highest success rate or biodiversity gain, implementations should be founded on empirical evidence (Sutherland et al. 2004; Ferraro and Pattanayak 2006; Samways et al. 2020). Especially for insects— whose decline and biological value get widespread attention (Didham et al. 2020; Wagner et al. 2021)—scientific evidence on the effects of landscapes on functional connectivity and gene flow is still limited (Keyghobadi et al. 1999; Wilcock et al. 2007; Keller et al. 2012; Pérez-Espona et al. 2012; Watts et al. 2016; Haran et al. 2017; Trense et al. 2021).

An indirect way of measuring functional connectivity is through molecular genetic data. With landscape genetics methods, one can quantify the effects of landscape composition between habitat patches on gene flow between populations (Manel et al. 2003; Manel and Holderegger 2013; Balkenhol et al. 2015). Patterns of gene flow reflect realised or successful dispersal and depend on post-dispersal survival and reproduction (Spear et al. 2010). Landscape genetics methods allow for inferences regarding connectivity without the need to collect individual movement data. Collecting such field data—for example from telemetry or mark-recapture methods—requires intensive effort or is even unfeasible for many organisms, such as the majority of insects (Spear et al. 2010; Batsleer et al. 2020). Only some larger insect species have been employed with tracking devices, and even for these, the biases on the resulting measurements remain insufficiently considered (Batsleer et al. 2020). Both mark-recapture and telemetry rarely detect, and thus underestimate, long-distance dispersal (Ugelvig et al. 2012; Trense et al. 2021; De Ro et al. 2021). The advancing tools in molecular and landscape genetics, such as in landscape resistance modelling, can help us understand connectivity and landscape use in species for which direct estimates of habitat use and movement are difficult to obtain with more classical field methods (Spear et al. 2010). However, such methods are rarely applied to insects. Studies on a few specialist insects point towards non-habitat and/or anthropogenic features forming barriers, such as in in the Sooty Copper butterfly *Lycaena tityrus* (Trense et al. 2021) and the army ant *Eciton burchellii* (Pérez-Espona et al. 2012). While for the long-horned beetle *Monochamus galloprovincialis*, which is considered an excellent disperser and even a vector for diseases in pinewoods (David et al. 2014), open areas do not seem a barrier to gene flow (Haran et al. 2017).

Coastal dunes in Flanders are geological young formations with calcareous sandy soils forming a narrow, elongated ecosystem along the coast (Provoost et al. 2004; Decleer 2007). They harbour a threatened, specific insect biodiversity typical of sandy early-succession and pioneer vegetations (Maes and Bonte 2006). In northwest Europe, and more specifically Flanders (Belgium), coastal dunes have gone through extensive landscape changes and fragmentation during the past century (Provoost et al. 2011). The fragmentation of coastal sandy habitats had two main impacts: loss of to talarea and decrease in habit at quality of theremaining patches. Urbanisation from the interbellum period (1920-1940) onwards decreased the total dune area by half during the previous century and separated the major dune entities physically (Provoost and Bonte 2004). The resulting obstruction of wind and sand dynamics, combined with the loss of agricultural practices (such as grazing of livestock), stimulated succession and scrub development, decreasing the amount of open dune habitats (Provoost et al. 2011). Large herbivores have been introduced in many coastal dune reserves to revitalise dune dynamics as a substitute for natural wind dynamics, but have—due to trampling—a variable effect on local arthropod species (Bonte and Maes 2008; van Klink et al. 2015; Batsleer et al. 2022b). Altogether, coastal dunes in Flanders are a human-altered landscape with complex nature management considerations to consider in both landscape scale and site-based nature management. In such a landscape, understanding functional connectivity—with its many specialist dune species—is vital to make evidence-based conservation decisions.

Conservation and connectivity studies have historically focused on plants and vertebrates, with insects receiving less attention. When insects are considered, the focus is typically on butterflies and pollinators (Clark and May 2002; Potts et al. 2016). Even for pollinators, the importance of nesting resources is often overlooked in nature management (Kimoto et al. 2012; Buckles and Harmon-Threatt 2019). As a result, ground-nesting Hymenoptera remain particularly neglected in both conservation and connectivity studies. The digger wasp *Bembix rostrata* is considered an emblematic representative of this group, associated with open and early-succession dune vegetations in Belgium (Batsleer et al. 2022b).

Here, we report on the genetic (functional) connectivity of the dune-habitat specialist digger wasp *B. rostrata* in a human-altered coastal landscape in Belgium using landscape genetics methods. While it is known from an earlier population genetic study that the species’ regional connectivity is large (Batsleer et al. 2024), it remains unknown which specific landscape components facilitate or constrain gene flow. Given the strong urbanisation of the coastal dunes, we expect this land-use to form a barrier to gene flow in this dune specialist insect. Given the species biology, we expect the dune areas and beaches to function as facilitators to gene flow. These hypotheses are parallel to results for other specialist insects in landscape genetics studies (Pérez-Espona et al. 2012; Trense et al. 2021). We use a resistance raster approach to quantify resistance values of the distinctive landscape types in our study area. The resistance values represent the degree to which each landscape type impedes genetic connectivity for the study species (Spear et al. 2010). To determine the resistance values of each landscape type, these values are optimised, by maximising the correlation between the landscape resistance distances and observed genetic distances between all pairwise individuals (Peterman 2018). All possible combinations of landscape categories were optimised to be able to perform multi-model inference. This allows us to deduce possible barriers or facilitators in the landscape for this dune-specialist digger wasp. Based on this estimate of functional connectivity, we can make recommendations on the balance between landscape- and site-based conservation.

## Material & Methods

### Study species

*Bembix rostrata* (Linnaeus, 1785; Hymenoptera, Crabronidae, Bembicinae) is a univoltine dune-specialist, gregariously nesting digger wasp from sandy habitats with sparse vegetation, sensitive to grazing management (Larsson 1986; Klein and Lefeber 2004; Bonte 2005; Batsleer et al. 2022b).

Adults are active in summer, showing protandry: females are immediately mated upon emergence by the guarding males, who emerge one to five days earlier (Wiklund and Fagerström 1977; Schöne and Tengö 1981; Evans and O’Neill 2007). Females show brood care: one individual constructs one nest burrow at a time in which it progressively provisions a single larva with flies (Nielsen 1945; Field et al. 2020). It is estimated that up to 5 nests (in northern regions) are produced, each with one offspring (Larsson and Tengö 1989). Nesting locations of *B. rostrata* usually span a width of less than 100 m, with nests arranged in several clusters that have a maximum diameter of 10 m (Batsleer et al. 2022a). Within these clusters, nest densities typically reach up to 10 nests per square meter (Nielsen 1945; Larsson 1986). There are no overlapping generations, as the species overwinters as prepupa and there is only one generation per year. *Bembix rostrata* is often considered to be a philopatric species, as they do not easily colonise vacant habitat and exhibit site fidelity throughout the nesting season (Nielsen 1945; Larsson 1986; Bogusch et al. 2021; Batsleer et al. 2022a). However, a population genetics study across Belgium showed that this does not preclude gene flow between existing populations, including some dispersal to distant, more isolated populations (Batsleer et al. 2024). This population genetics study showed that *B. rostrata* has relatively high genetic connectivity in the coastal dunes in Belgium: genetic distances were low, genetic compositions between populations substantially overlapped, a large number of immigrant links between coastal population were detected and an overall pattern of isolation-by-distance was observed (Batsleer et al. 2024).

### Genetic sampling

Sampling took place in the summer of 2018 in the dunes along the Belgian west coast (Fig. 1), from the French-Belgian border (De Panne) to the Yser estuary (Nieuwpoort), stretching 14 km along the coast and reaching maximum 4 km inland. As nests are clustered together at nesting aggregates in grey dunes (EU Habitats Directive habitat 2130), we tried to spatially cover all known nesting locations in the landscape and locally took samples spatially spread within a nesting location, across clusters of nests. Sample locations were based on a spatially exhaustive survey of potential nesting sites the previous summer. The coordinates of each individual sample were saved. Only females were sampled, to solely use diploid individuals in the genetic analyses for this haplodiploid mating system (Zayed 2009; Batsleer et al. 2024). To minimise the impact of sampling, we used a non-lethal sampling procedure with wing clips, a method shown to produce good-quality DNA for microsatellite PCR amplification and having no clear behavioural effects for honey bee queens and workers (Châline et al. 2004). Such wing clips of less than 10% of the wing area mimic natural wear and are shown not to have an effect on metabolic flight costs in bumblebees (Hedenström et al. 2001). Forewing tips from live digger wasps were cut on both sides and stored in absolute ethanol, kept at 4°C after each day of sampling and transferred to a freezer (−18°C) for longer term storage. The individual digger wasps displayed normal behaviour when flying off after sampling.

**Figure 1:**
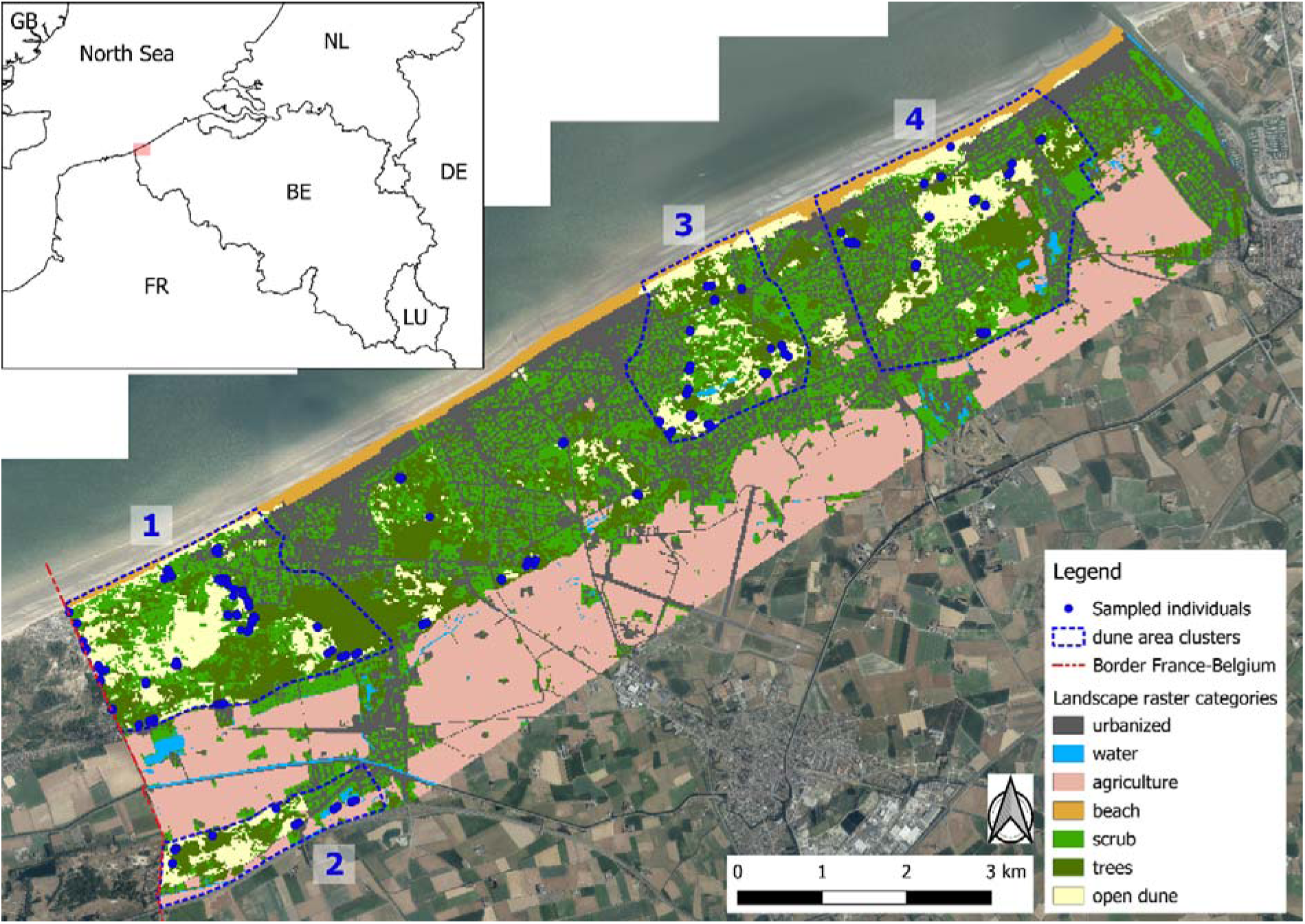
overview map of study area: west coast in Belgium from the French-Belgian border (west; dashed red line) to the Yser estuary (east). Sample locations of each individual (blue dots) are shown and four dune area clusters (blue dashed lines) are shown. Four dune area clusters (blue dashed lines) are ecological clusters deemed relevant to nature management; 1) Westhoek 2) Cabour 3) Doornpanne 4) Ter Yde. The landscape raster categories are shown with a 20m × 20m resolution (cell size). Neighbouring countries of Belgium (BE): France (FR), Luxembourg (LU), Germany (DE), The Netherlands (NL), United Kingdom (GB). Map made with QGIS v3.22 (QGIS Development Team 2020) in Cartesian coordinate reference system Belgian Lambert 72, EPSG:31370. Background aerial photographs (summer 2018) from Agency for Information Flanders (source: geopunt.be).

DNA extraction and PCR amplification were performed as described in Batsleer et al. (2024) and summarised hereafter. DNA is extracted from the wing tips with a protocol based on Chelex (Biorad). 33 polymorphic microsatellites are PCR-amplified for each sampled individual and electropherograms are scored using Geneious Prime 2021.0 (Biomatters). Microsatellites that had a lot of stutter in amplification (five microsatellites) or that were excluded after assumption testing (Hardy-Weinberg, null alleles and linkage disequilibrium; seven microsatellites) in the population genetics study in (Batsleer et al. 2024) were also excluded here, resulting in 21 microsatellites (out of 33) for the subsequent genetic analysis. If an individual had more than 8 missing loci, it was discarded from the analyses (2.4%, 13 out of 552). A total of 539 sampled individuals were successfully amplified. The resulting data contained an overall of 0.7% missing data, which were replaced by the mean allele frequencies for the calculation of the genetic distances (see further).

### Land cover categories

We combined two types of land cover datasets available for the study area. For landscape categories within dune areas, we used detailed vegetation maps (resolution 0.4m) available for the region derived from remote sensing data with a classification machine learning technique (Bonte et al. 2021). For landscape categories outside dune areas, we used the ground cover map 2018 provided by the Flemish government (‘Bodembedekkingskaart’ BBK 1m resolution, source: www.geopunt.be). We aligned, reclassified and combined the raster datasets (see online resource 1 for detailed information on reclassificaitons) to have one raster file with resolution of 1m of the study area with 7 simplified landscape categories: urbanised*u*(*rban*), *water*, agricultural (*agric*), *beach*, *scrub*, *trees* and open dune (*opend*) vegetation (Fig. 1). Roads and urban features were analysed as a single category, as roads are almost always embedded or aligned within urban area, resulting in a high degree of spatial dependency. Category *beach* was added by rasterizing a manually drawn shapefile based on aerial maps and the high water line. For further analyses, we resampled the rasters to work with resolutions of 20 m and 50 m. We deem these resolutions (spatial grains) adequate to capture the turnover in heterogeneity of the focal landscape and to be linked with the ecological processes investigated. When relevant landscape elements are captured by the resolution, landscape genetic analyses should be relatively robust for changes in cell size (McRae et al. 2008; Cushman and Landguth 2010).

### Genetic distance measure

We used individual-based genetic distances between pairwise individuals. We used a metric based on multiple axes of a principle components analysis (PCA), which was shown to perform better in landscape genetics analyses than other individual-based metrics, especially when sample size and genetic structure are low (Shirk et al. 2017; Kimmig et al. 2020). First, principal components (PC) were calculated from a multiple contingency table (0, 1, 2 values for each allele for each individual).

Second, the Euclidean distance based on the first 16 PCs was calculated to create a genetic distance matrix between all individuals (Shirk et al. 2017). To calculate these genetic distances, we used the R-packages adegenet v2.1.10 (Jombart 2008) and ecodist v2.1.3 (Goslee and Urban 2007).

### Landscape resistance optimisation

The unknown values are the resistance values of each landscape type, which quantify at each grid cell the constrain to gene flow (Spear et al. 2010). Based on these resistance values, resistance distances between pairwise individuals are iteratively recalculated during the optimisation (see further for details). Such resistance distances reflect a measure of separation between pairwise individuals and incorporate the effects of different permeabilities of the landscape (Spear et al. 2010). The resistance values are optimised based on maximising the correlation between genetic distances and landscape resistance distances between pairwise individuals. Thus, it is expected that the effect of the landscape on genetic connectivity has a detectable signal on the observed genetic distances. The optimisation was implemented with the RESISTANCEGA R package v4.0-0 (Peterman 2018).

Resistance distance measure. We used the commute resistance distance in RESISTANCEGA (van Etten 2017; Peterman 2018), which is the expected time of a random walk (the stochastic process of random steps) between a pair of points (sampled individuals) along a circuit-based resistance surface (the raster with resistance values). This type of distance accounts for several possible pathways through the landscape (McRae et al. 2008; Peterman 2018). The pairwise resistance distances are recalculated at each iteration based on the resistance values of the landscape types during the previous iteration (with initial values of 1).

Optimisation procedure. RESISTANCEGA uses a linear mixed effects model with maximum likelihood population effects (MLPE) parameterisation to fit pairwise resistance distances (predictor) to pairwise genetic distances (response). MLPE parameterisation accounts for the non-independence among pairwise data (Clarke et al. 2002; Bates et al. 2015). The optimisation procedure iteratively searches many possible resistance value combinations to maximise the MLPE model fit (based on log-likelihood), resulting in the best fit between resistance distances and genetic distances (Peterman 2018). This optimisation procedure is based on a genetic algorithm (Scrucca 2013). For each raster, we used the SS_OPTIM function in R ESISTANCEGA to optimise the resistance values (see online code).

Parameters were set to default, only the maximum number of iterations (*max.iter*) was changed to 300. There are no a priori expectations for the resistance values of the landscape categories, as no information is present regarding habitat selection and movement behaviour at the landscape scale for this species (Knowlton and Graham 2010). Consequently, and by default, the resistance values of each category have an initial value of 1. Multicategorical rasters. We optimised multi-categorical rasters, which consider several landscape categories at once. Although more complex to parameterise, multi-categorical raster optimisation adds more biological realism to the analysis than considering only one single category at a time, which would assume an organism’s perception of a landscape to be binary (Spear et al. 2010). We ran optimisations for each possible combination of all 7 categories. A raster then consists of one to seven focal landscape categories and all else or ‘the rest’ (for example when categories *urban* and *scrub* are considered, all five other categories are lumped into a third rest category). The total number of all possible combinations of 7 categories (*urban*, *water*, *agric*, *beach*, *scrub*, *trees* and *opend* vegetation) is 127: Σ^7^_k=0_(7 k). Ascii-files needed as input for RESISTANCEGA were created for each possible combination with the R-package terra v1.7.78 (Hijmans 2024), see online code. Thus, optimisations are run for each possible combination for the 7 categories.

Bootstrap analysis. To compare the performance of each of the 127 optimised rasters and the Isolation-By-Distance (IBD) as null model, we performed a bootstrap analysis. With the RESIST.BOOT function from R ESISTANCEGA, 75% (default) of the individuals are subsampled and the MPLE model refitted for each of the 127 optimised rasters and IBD, this for 1000 iterations. Relative support for each model is then compared using several bootstrap-derived metrics for model performance, importance and stability. The primary metric used is the difference in corrected Akaike information criterion (AICc, correction for small sample sizes; measure for predictive accuracy of a model). In general, it is considered that models with ΔAICc < 2 are competing models, and even models with ΔAICc ranging between 2-7 have some support and should not be dismissed (Burnham et al. 2011).

Other metrics included are: number of times the model was the top model (n.top), the average rank (avg.rank, measure for overall relative importance), and the average Akaike weight (avg.weight, measure for relative contribution), average marginal R² (explained variance) and log-likelihood (predictive fit) across bootstrap samples. The extra bootstrap-metrics indicate, next to ΔAICc, if certain models are estimated to be consistently more important than other models across bootstrap samples.

Independent optimisations runs. As the optimisation with R ESISTANCEGA is a stochastic process, optimised values can differ from run to run, and it is advised to run all optimisations at least twice to confirm convergence and relative relationships among resistance values (Peterman 2018). To check this convergence and account for a possible bias due to the starting cell values of a category in multi-categorical rasters (i.e. the initial order: urban, water, agricultural, beach, scrub, trees, open dune) (Peterman 2018; Kimmig et al. 2020), we ran the multi-categorical raster optimisations and bootstrap analysis a second time with the order of categories inverted (starting with *opend*, ending with *urban*). To confirm convergence more conclusively, we ran optimisations and bootstrap analyses of the seven single category rasters (that consider only one category) four times each (online resource 2). This is computationally less intensive than running all possible combinations, but gives a broad idea of the robustness of the optimisation.

Multi-model inference. As all possible combinations of categories were optimised to obtain resistance values, we were able to use a multi-model inference approach to obtain robust estimates of average weight for each category and average relative resistance value. Multi-model inference is a procedure to account for the uncertainty of parameter estimates related to model selection and is especially useful when no single model is overwhelmingly supported by the data (Johnson and Omland 2004). We used Akaike weights from the bootstrap analysis considering all the models (i.e. optimised rasters) for each area type. From these, we calculated the summed Akaike weight per category by taking into account a model if the focal category was included (Burnham and Anderson 2004; Johnson and Omland 2004). These summed Akaike weights give an indication for the relative importance of each category to explain the pairwise genetic distances. We also calculated relative resistance values of each category. As optimised resistance values depend on the other categories included in a multi-categorical model, interpretation for a focal category should be done relative to these other categories. To be able to do this for each landscape category, we rescaled the resistance values per optimised raster: we subtract the resistance value of the focal category from the other resistance values that were simultaneously optimised. As such, the value of the focal category becomes zero and is considered facilitating to gene flow compared to categories with values larger than zero and a barrier to gene flow compared to categories with values smaller than zero (for further information on interpretation, see results section). All models and the relative resistance values per focal category were then summarised by plotting boxplots for each focal category with ggplot2 v3.5.1 (Wickham 2016). We consider this as a qualitative type of model averaging in multi-model inference, where parameters are summarised across all models (Johnson and Omland 2004).

We remain descriptive with these relative resistance values, as values used in these boxplots are not independent because categories are optimised simultaneously.

To get an idea of the possible small scale variation in landscape effects on gene flow, we repeated the analyses for each of the four dune area clusters within our study area (Fig. 1). These dune areas are seen as ecological clusters relevant for nature management as these are clusters of the largest local nature reserves (Fig. 1). These optimisations for the dune area clusters are run with a resolution of 20 m (in contrast to 50 m for the complete study area). Details and results can be found in online resource 4.

The total computational time (runs for all possible combinations of categories) was very high and could not be run on a single desktop. Therefore, we made use of the high-performance computing infrastructure of Ghent University (HPC-UGent) to run the optimisations and bootstrapping analyses. By doing this, we could run all possible combinations in the multi-categorical raster optimisations, which is the most ideal for robust statistical inference and allows for model averaging to assess the importance of the different landscape categories. Previous studies reported computational constraints for optimisations with RESISTANCEGA considering all combinations and used alternative approaches for model comparison based on model selection (e.g. Lourenço et al. 2019; Kimmig et al. 2020).

The best-supported models (i.e. optimised rasters) for the complete study area were used to visualise the putative gene flow currents in the landscape using Circuitscape (McRae et al. 2013; Kimmig et al. 2020). Gene flow currents in Circuitscape represent the putative genetic connectivity in a landscape, represented by conductance values for a grid cell, where most gene flow passes between sample locations in the given landscape. Current maps were inferred for each pair of sample locations (representing multiple possible pathways in the landscape). The conductances of these current maps were all summed for the final Circuitscape current map, to give a representation of the overall genetic connectivity of the landscape (McRae et al. 2008; Kimmig et al. 2020).

The main analyses (see online code) were performed in RS TUDIO V1.4 (RStudio Team 2022) using R v4.1.3 (R Core Team 2020), except for calculations for the current map with the GUI of Circuitscape v.4.0.5 (McRae et al. 2013) and cartographic visualisations in QGIS V3.22 (QGIS Development Team 2020).

## Results

Below, we first describe the results of the optimisation of the resistance values of the multi-categorical landscape maps after bootstrapping. The results show which optimised landscape resistance models relatively perform best to explain the gene flow of *Bembix rostrata* in the study area. The optimisations provide which landscape categories are facilitating or forming a barrier to gene flow for each landscape model. Second, we give results on the multi-model inference to deduce overall patterns of landscape categories across all optimised models. The results indicate which landscape categories are, among all optimised landscape models, the most important in explaining the gene flow and what their relative resistance values are. We further provide a visualisation of Circuitscape connectivity maps of the three best-supported models and describe the results of the analyses on the smaller scale dune area clusters within the study area.

### Bootstrap analysis of landscape resistance optimisation

Results of the top five best supported models with optimisations of multi-categorical models (all possible combinations of landscape categories) after bootstrapping can be found in table 1 (full table in online resource 3). In general, there was no overwhelming support for one or a few models based on ΔAICc, as isolation-by-distance (null model) was one of the best supported models (Distance; ΔAICc=1.20 run 1 and 1.42 run 2; table 1) and was a competing model considering ΔAICc < 2 with *urban* (the best supported model). In general, R²m’s were low (table 1; overall <1%), Akaike weights were small (avg.weight in table 1) and all ΔAICc’s were relatively small (table 1; maximum ΔAICc was 14.58; online resource 3), and even models with ΔAICc ranging between 2-7 have some support and should not be dismissed (Burnham et al. 2011).

**Table 1:**
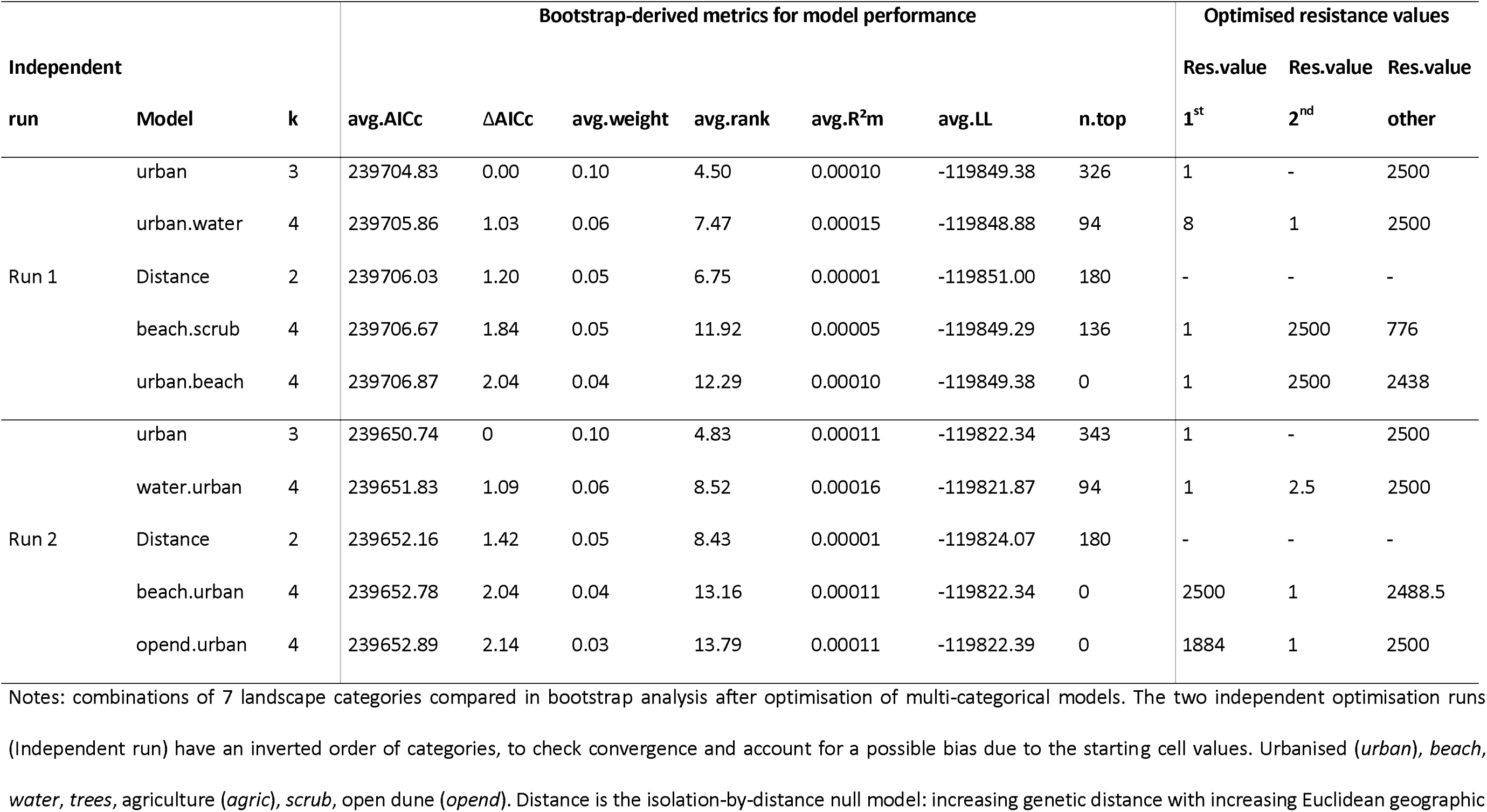
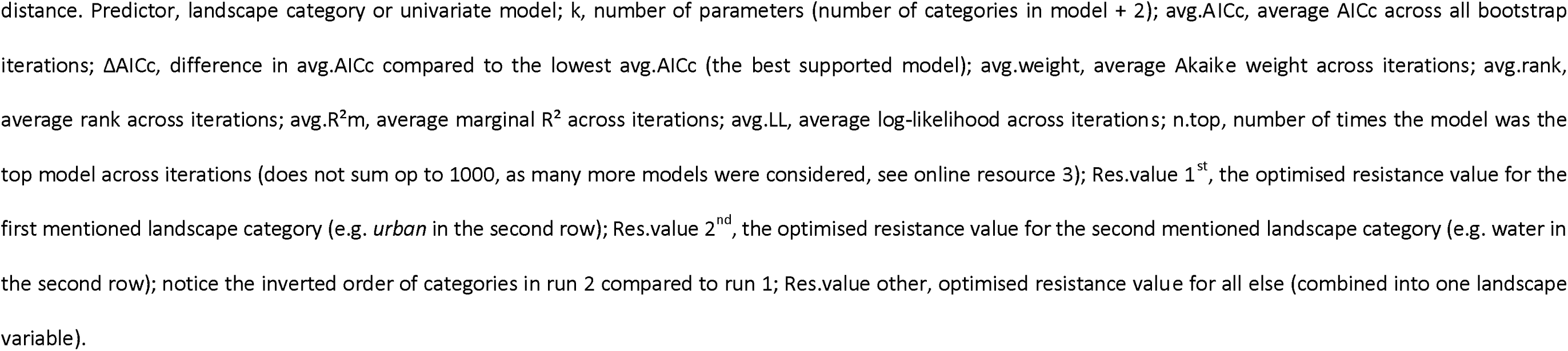
Top five bootstrap results for the multi-categorical model optimisations for the complete study area for the two independent optimisation runs. Full table with all models (and null model) can be found in online resource 3.

| Independent |  |  | Bootstrap-derived metrics for model performance |  |  |  |  |  |  | Optimised resistance values |  |  |
| --- | --- | --- | --- | --- | --- | --- | --- | --- | --- | --- | --- | --- |
| | | | avg. AICc | ΔAICc | avg. weight | avg. rank | avg. R<sup>2</sup>m | avg. LL | n.top | Res.value | Res.value | Res.value |
| run | Model | k |  |  |  |  |  |  |  | 1<sup>st</sup> | 2<sup>nd</sup> | other |
| Run 1 | urban | 3 | 239704.83 | 0.00 | 0.10 | 4.50 | 0.00010 | -119849.38 | 326 | 1 | - | 2500 |
|  | urban.water | 4 | 239705.86 | 1.03 | 0.06 | 7.47 | 0.00015 | -119848.88 | 94 | 8 | 1 | 2500 |
|  | Distance | 2 | 239706.03 | 1.20 | 0.05 | 6.75 | 0.00001 | -119851.00 | 180 | - | - | - |
|  | beach.scrub | 4 | 239706.67 | 1.84 | 0.05 | 11.92 | 0.00005 | -119849.29 | 136 | 1 | 2500 | 776 |
|  | urban.beach | 4 | 239706.87 | 2.04 | 0.04 | 12.29 | 0.00010 | -119849.38 | 0 | 1 | 2500 | 2438 |
| Run 2 | urban | 3 | 239650.74 | 0 | 0.10 | 4.83 | 0.00011 | -119822.34 | 343 | 1 | - | 2500 |
|  | water.urban | 4 | 239651.83 | 1.09 | 0.06 | 8.52 | 0.00016 | -119821.87 | 94 | 1 | 2.5 | 2500 |
|  | Distance | 2 | 239652.16 | 1.42 | 0.05 | 8.43 | 0.00001 | -119824.07 | 180 | - | - | - |
|  | beach.urban | 4 | 239652.78 | 2.04 | 0.04 | 13.16 | 0.00011 | -119822.34 | 0 | 2500 | 1 | 2488.5 |
|  | opend.urban | 4 | 239652.89 | 2.14 | 0.03 | 13.79 | 0.00011 | -119822.39 | 0 | 1884 | 1 | 2500 |

Notes: combinations of 7 landscape categories compared in bootstrap analysis after optimisation of multi-categorical models. The two independent optimisation runs (Independent run) have an inverted order of categories, to check convergence and account for a possible bias due to the starting cell values. Urbanised (*urban*), *beach*, *water*, *trees*, agriculture (*agric*), *scrub*, open dune (*opend*). Distance is the isolation-by-distance null model: increasing genetic distance with increasing Euclidean geographic
distance. Predictor, landscape category or univariate model; k, number of parameters (number of categories in model + 2); avg.AICc, average AICc across all bootstrap 445 iterations; ΔAICc, difference in avg.AICc compared to the lowest avg.AICc (the best supported model); avg.weight, average Akaike weight across iterations; avg.rank, average rank across iterations; avg.R<sup>2</sup>m, average marginal R<sup>2</sup> across iterations; avg.LL, average log-likelihood across iterations; n.top, number of times the model was the top model across iterations (does not sum up to 1000, as many more models were considered, see online resource 3); Res.value 1<sup>st</sup>, the optimised resistance value for the first mentioned landscape category (e.g. *urban* in the second row); Res.value 2<sup>nd</sup>, the optimised resistance value for the second mentioned landscape category (e.g. water in the second row); notice the inverted order of categories in run 2 compared to run 1; Res.value other, optimised resistance value for all else (combined into one landscape variable).

### Multi-model inference

When no single model is overwhelmingly supported by the data, multi-model inference is ideal to deduce average weights and relative resistance values of categories across all models (Johnson and Omland 2004). From such a multi-model inference based on the optimisation of all possible combinations, we deduced more general patterns from the data based on the summed Akaike weights per landscape category (table 2) and the relative resistance values (Fig. 2). *Urban* was an important explanatory category to explain pairwise genetic distances (table 2), which acted mainly as facilitator to gene flow compared to other landscape categories (table 1, Fig. 2). Natural landscape categories (*beach*, *scrub*, *trees*, *open dune*; bottom row in Fig. 2) were in general more resistant to gene flow than anthropogenic categories (*urban*, *agriculture, water*; top row in Fig. 2) in the focal landscape. As most categories, apart from urban, were of little (and inconclusive) importance to explain pairwise genetic distances (table 2), the uncertainty of relative resistance values between these categories (especially natural categories, Fig. 2 second row) should be considered high. The second independent optimisation run (included in table 1 and 2; Fig. 2) confirmed these patterns and showed that category *urban* was consistently present in the top best supported models (table 1) and the most important (and only conclusive) category to explain pairwise genetic distances (table 2).

**Figure 2:**
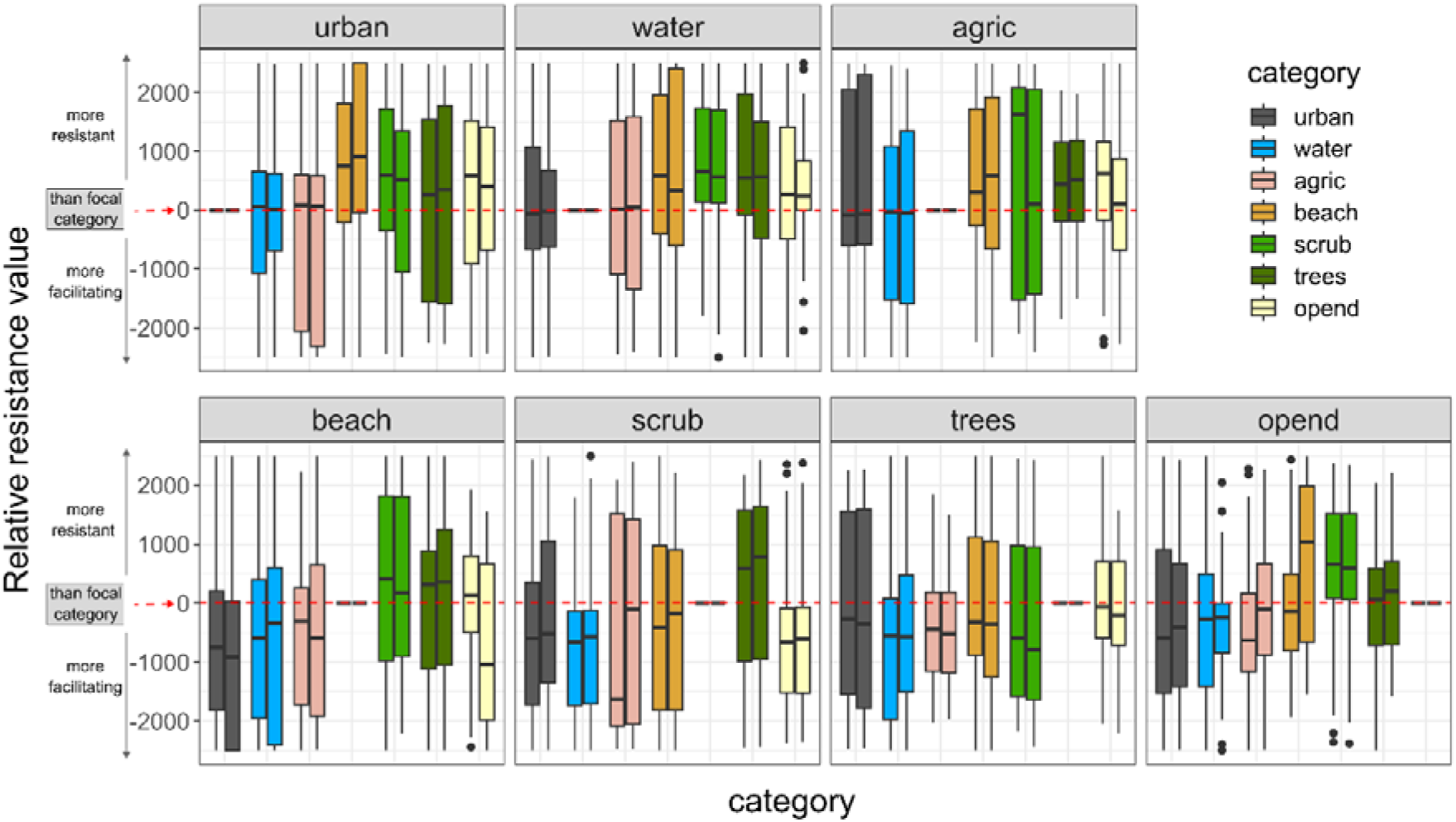
Relative resistance values from the optimisation of multi-categorical models for the complete study area, as part of the multi-model inference. Anthropogenic landscape categories (urban, water, agric=agriculture) are in the top row, natural landscape categories (beach, scrub, trees, opend=open dune) in the bottom row. The two independent runs are combined as paired boxplots for each category in each panel (left run 1, right run 2). Resistance values are rescaled for each focal category: as optimised resistance values depend on the other categories included in a model, all models that include the focal category are selected. The focal category is then held constant by subtracting its optimised resistance value in each model from the optimised resistance values of the other models (resistance values - resistance value of focal category). These rescaled values are then plotted in boxplots for each focal category. Each panel should then be interpreted by checking what lies above zero (category more resistant than focal category) and below zero (category more facilitating than focal category). The red dashed line is zero, the boundary of equal resistance values between the focal category (panel) and other categories. For example, in the urban panel (left upper corner), the medians of beach, scrub, trees and open dune are situated above the red line, consequently, those four categories are in general more resistant than urban (urban is more facilitating) to gene flow. As most categories, apart from urban, proved to be of little importance to explain pairwise genetic distances (table 2), the uncertainty on the relative resistance values of these categories should be considered high.

**Table 2:** Summed Akaike weights per category for multi-categorical model optimisations, as part of the multi-model inference. The sum of the weights indicates the importance of a category in all possible multi-categorical optimised rasters to explain the pairwise genetic distances.

| Category | Sum weights run 1 | Sum weights run 2 |
| --- | --- | --- |
| urban | 0.496 | 0.501 |
| agric | 0.357 | 0.335 |
| beach | 0.356 | 0.376 |
| trees | 0.338 | 0.307 |
| water | 0.297 | 0.315 |
| scrub | 0.286 | 0.259 |
| opend | 0.246 | 0.258 |

Notes: the Akaike weight for an optimised model was added to the summation (Sum weights) if the focal category (Category) was in it.

This was also confirmed by the four independent runs of the single categories, which consistently had urban as the first supported model and IBD as the second best competing model (online resource 2).

### Visualisation current maps

Circuitscape current (or flow) maps (Fig. 3) were made for the three best supported models (table 1). As all ΔAICc’s were relatively small (online resource 3), there were many more models worth considering (as was done in the multi-model inference, Fig. 2 and table 2). However, these Circuitscape current maps give post-hoc visualisations of how the landscape likely influences genetic connectivity within the study area according to the best three models.

**Figure 3:**
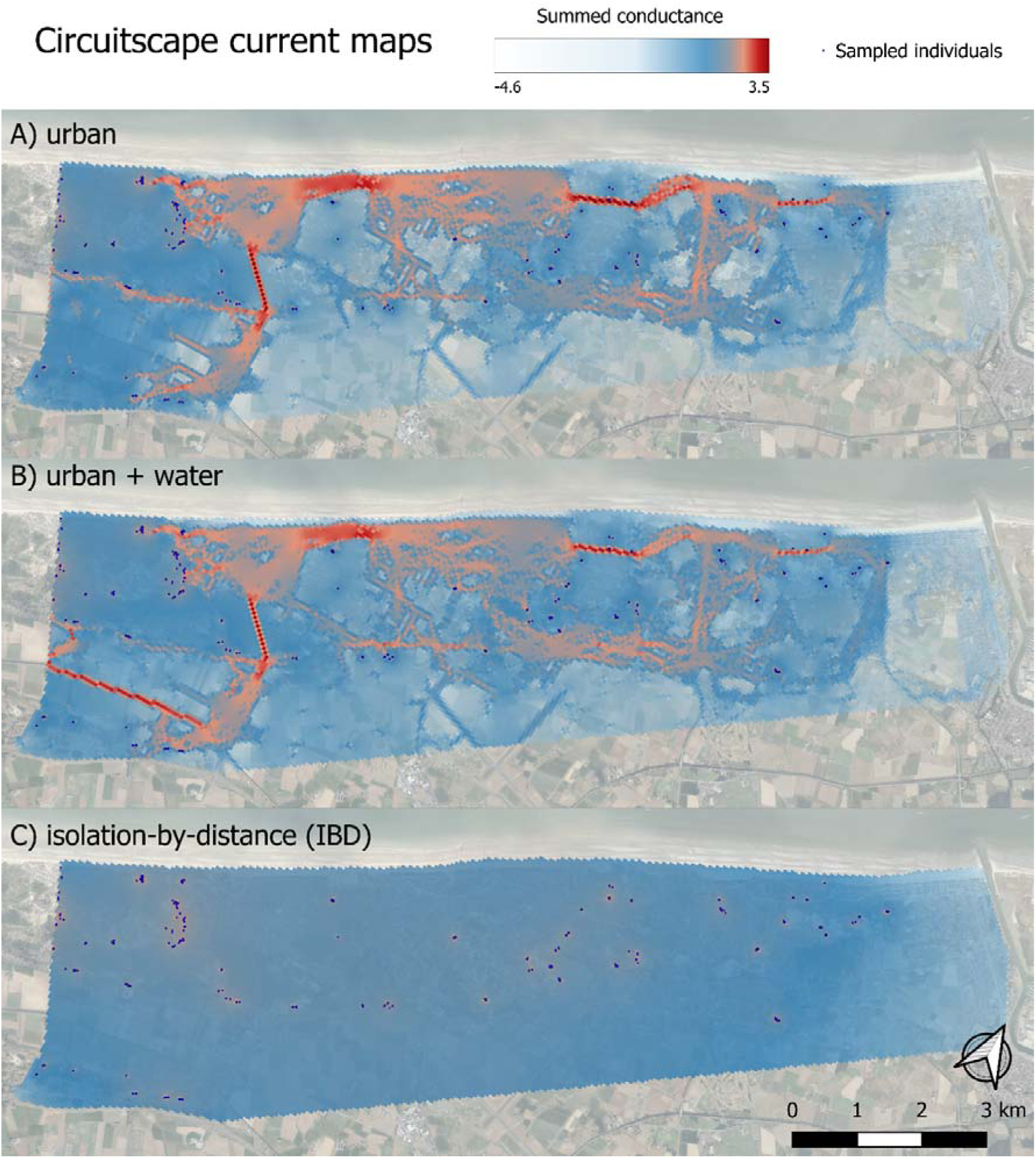
putative genetic connectivity with CIRCUITSCAPE current maps for the complete study area for the three best supported models. These maps represent in which grid cells of the landscape the most gene flow passes between the sampled nesting locations (black dots), based on A) best supported model with urban as facilitator, B) 2 best supported model urban and water combined, both facilitators and C) 3 best supported model Isolation-by-distance (resistance values for all cells was set to 1). The values of the grid cells are log10-transformed summed conductance (a relative measure with no unit), which is based on the summation of all current maps between pairwise sampled individuals (black dots). High conductance in dark red, medium conductance in blue, low conductance in white-transparent. A map with the corresponding land cover categories can be found in figure 1. Map made with QGIS v3.22 (QGIS Development Team 2020). Background aerial photographs (summer 2018) from Agency for Information Flanders (source: geopunt.be).

### Small-scale dune area clusters

The analysis repeated at the level of each of the four dune area clusters (Fig. 1) showed Isolation-By-Distance (IBD) to be the overall best supported model on this smaller scale (online resources 4 and 5). Best supported models and the multi-model inference showed some smaller variation among the studied clusters (online resource 4) and generally confirmed the overall pattern from the complete study area: anthropogenic categories (*urban*, *agric*, *water*) were more facilitating to gene flow than natural landscape categories. There were two exceptions: for the dune area Doornpanne (dune cluster 3 in Fig. 1), *trees* was a facilitating landscape property to gene flow, and for Cabour (dune cluster 2 in Fig. 1), *agriculture* showed to be a strong barrier to gene flow. This is probably related to the specific habitat composition and landscape context for these dune areas: Doornpanne has relatively much encroached area (trees and shrubs), while Cabour is the dune area most embedded in agricultural land (Fig. 1).

## Discussion

We used landscape genetics to unravel patterns of functional connectivity of the digger wasp *Bembix rostrata* in a human-altered coastal region. Despite the species’ strong gene flow at regional scales and overall pattern of isolation-by-distance (Batsleer et al. 2024), we were able to discern signals of the landscape on the genetic patterns within the coastal dune metapopulation. Contrary to expectations, we found a consistent signal of urban features facilitating gene flow. In general, anthropogenic landscape categories (urban, agriculture, water) seem more facilitating than natural habitat types in this human-altered dune landscape context.

In line with the previous study at larger spatial scales (Batsleer et al. 2024), isolation-by-distance (IBD) remained a prominent pattern of the species’ landscape genetics. In itself, an IBD pattern is not in contradiction with an overall high genetic connectivity or an effect of the landscape (Keyghobadi et al. 1999). An IBD pattern emerges naturally when dispersal is limited at a certain spatial scale, i.e. dispersal is more likely to nearby populations than distant ones, independent of the landscape (Kimura and Weiss 1964). Consequently, IBD is seen as a biological realistic null model (opposed to panmictic gene flow) in landscape genetics when considering the influence of the landscape on gene flow, i.e. isolation-by-resistance (IBR) (Wagner and Fortin 2015). Our results showed that IBD was consistently a competing model and one of the best supported models (table 1, considering ΔAICc < 2 with *urban*). Because overall support for models was low (low values of ΔAICc’s), we applied multi-model inference to deduce average weights and relative resistance values of categories (Johnson and Omland 2004). This confirmed the importance of the urban category for discriminating pairwise genetic distances and deduce that generally anthropogenic landscapes seem more facilitating than natural habitat types. Therefore, considering all possible combinations for landscape resistance optimisation can be powerful when landscape resistance signals are weak and hard to detect.

The current maps in figure 3 visualise the putative realised conductance of grid cells based on all sample locations according to the three best supported and competing models (*urban*, *urban*-*water*, IBD; table 1). These current maps are derived from the optimised resistance raster and serve as post hoc visualisations of the inferred relationship between genetic distance and resistance distance. In this context, the few isolated stretches of road with exceptionally high conductance values (Fig. 3B, 3C) are likely artifacts of this post hoc approach, resulting in overly inflated values. Therefore, the primary focus should be on the overall patterns. In general, we can see from these current maps that the best-supported models should be considered as complementary rather than opposing. First, waterways often run parallel to roads and waterbodies are embedded along or within urban areas (Fig. 1). Such water—which is also a scarce landscape type—adds only a small extra effect on top of the urban areas for the resulting current map (Fig. 3B). Consequently, we cannot easily disentangle the effect of water from that of urban within the given landscape (Angelone et al. 2011; Keller et al. 2012). The Akaike weights from multi-model inference (table 2) indicate that the urban category is overall the most important to explain pairwise genetic distances. Second, the IBD pattern (Fig. 3C) is also present within the dune areas in the current map for the *urban* and *urban*-*water* models (high conductance around sampled individuals; Fig. 3A, 3B). In the latter models, the natural habitat categories (*open dune*, *scrub*, *trees*, *beach*) all have the same resistant values (relatively higher than urban and/or water; table 1), resulting in a homogenous resistance landscape within dunes, resulting in, on average, an IBD pattern. The importance of IBD on this scale is also confirmed by the analyses repeated on the dune area clusters within our study area (online resource 4). This interpretation is also in agreement with the multi-model inference, as natural habitat categories proved to be of little importance to explain pairwise genetic distances (table 2) and the uncertainty on the relative resistance values of these categories was high (Fig. 2). Consequently, IBD can be considered as the main background process, wherein gene flow depends on the pairwise, straight-line distances. The extra signal from anthropogenic features is then at play between dune areas where especially urban areas give a weak but consistent extra signal of conductance.

Contrary to our hypothesis, we found that gene flow of a specialised digger wasp nesting in grey dune habitat was not facilitated by typical beach and dune habitats, but rather by urban features in a dune landscape. This finding does not align with other landscape genetics studies on specialist insects, for which non-habitat landscape types were mostly barriers (Pérez-Espona et al. 2012; Trense et al. 2021). Importantly, urban features, although found to be facilitating to gene flow, should not be interpreted as being favourable for this specialist dune insect. Patterns of gene flow only reflect a certain component of movement in a species and interpretation of results in a larger ecological framework should consider other crucial components of a species survival and resource requirements (Spear et al. 2010; Cushman and Lewis 2010). As *B. rostrata* is a dune-specialist insect, urban areas should be considered as inhospitable environments. As the deduced resistance values per category are relative values (Fig. 2), the pattern could be explained by both slower, confined movement in dune habitat and/or faster, compensatory movement in inhospitable areas. Indeed, search behaviour in unfamiliar environments can be more directional and faster than in natural, familiar habitats, where movements are more exploratory (Van Dyck and Baguette 2005; Schtickzelle et al. 2006; Knowlton and Graham 2010). Suitable habitat intervening the landscape might hinder gene flow between more distant populations, as individuals are likely to settle at encountered habitat during dispersal (Adriaensen et al. 2003; McRae et al. 2008; Keller et al. 2012). On the other hand, compensatory movement through inhospitable habitats can result from a lack of resources or can be a strategy to reduce mortality risk (Schtickzelle et al. 2006; Peterman et al. 2014). Both can be counterbalanced by moving faster in inhospitable environments, resulting in increased genetic connectivity between fragmented habitats (Schtickzelle et al. 2006).

The deduced resistance values are values per grid cell and relative compared to the other considered landscape types. Consequently, the deduced resistance values of a category will depend on the habitat composition and the spatial configuration of how populations are embedded within the landscape (Richardson et al. 2016; Haran et al. 2017). In our study, urbanisation induces large-scale fragmentation compared to more natural processes like shrub and woodland encroachment. Hence, a low resistance of urban infrastructure does not conflict with an increased isolation-by-distance in a more fragmented dune landscape, resulting in lower amounts of gene flow between isolated dune fragments. Differences in a species’ ecology (and evolutionary history) across regions could further complicate and diversify possible responses to a certain landscape type in different landscapes, especially at larger scales and across regions (Segelbacher et al. 2010; Spear et al. 2010). Therefore, results regarding the influence of anthropogenic landscape types on the gene flow of *B. rostrata* cannot easily be extrapolated or generalised. Nevertheless, our results for this species will reasonably remain valid for similar landscapes at comparable scales and with similar compositions, such as several human-altered coastal landscapes of northwest Europe. But extrapolation to inland sandy regions, where the species occurs as well, would be invalid. There, the degree of isolation between habitat patches (Batsleer et al. 2024) might decrease the tendency of the species to cross inhospitable matrix, increasing the resistance values (Schtickzelle et al. 2006).

We find that the connectivity of *B. rostrata* does not seem limited by the landscape composition within dune clusters and between the different dune entities. Even on the contrary, on top of the general process of IBD, inhospitable urban features seem to facilitate gene flow in this landscape for a given distance between different habitats. This indicates that for *B. rostrata*’s persistence in this study area, landscape-based conservation measures aimed at increasing connectivity between dune areas (e.g. with habitat corridors within and/or peripheral to the urbanised regions) would not be an effective management measure unless they are able to give rise to the establishment of new populations. It is probably more critical to increase or maintain large population sizes, by increasing habitat area and habitat quality (Richardson et al. 2016; Watts et al. 2016). The effectiveness of increasing connectivity in this landscape might be worth more consideration for cursorial species experiencing high mortality costs when crossing urban infrastructure (e.g., natterjack toad, Cox et al. 2017). However, benefits gained from landscape-scale and site-based conservation should always be balanced to maximise biodiversity benefits with the limited resources at hand (Watts et al. 2016). For the conservation of *B. rostrata*, other processes linked with dune habitat fragmentation are anyway more fundamental to consider than landscape connectivity, such as the decrease in habitat area itself and shifts in ecological processes (e.g. edge effects and ‘ecosystem decay’ such as inhibition of sand dynamics in dunes; Pfeifer et al. 2017; Chase et al. 2020). This fragmentation due to urbanisation decreases the overall number of potentially suitable locations, increases isolation and affects regional levels of exchange among populations in a spatially altered population network (Cheptou et al. 2017). Since urban areas are not a barrier to direct gene flow, these more indirect effects of fragmentation should be the focus to secure persistence of *B. rostrata* in the human-altered dune landscape. In the contemporary context of dune nature management, a site-based conservation approach to maintain and improve open dune habitat is recommended, for which grazing with large herbivores (cattle, horses) is a crucial management tool. However, a reconciliation is needed between short-term negative effects of trampling by grazers on nest densities of *B. rostrata* and long-term positive effects of maintaining open dune vegetations within dune areas. Therefore, a local adaptive management approach with sheep grazing is recommended (Bonte 2005; Batsleer et al. 2022b). Additionally, improving habitat quality of open dune vegetations through landscape-scale conservation aimed at restoring and increasing sand dynamics—rather than connectivity—would be a larger scale, sustainable measure for the persistence of *B. rostrata* in the coastal dunes in Belgium.

## Statements & Declarations

## Acknowledgements

We thank the following persons and instances for permission and access to nature reserves: Johan Lamaire, Guy Vileyn, Koen Maertens, Evy Dewulf and Klaar Meulebrouck from ANB (Agency for Nature and Forests – Flemish government); Rika Driessens from IWVA/Aquaduin. We thank two anonymous reviewers for their constructive peer-review comments.

The computational resources (Stevin Supercomputer Infrastructure) and services used in this work were provided by the VSC (Flemish Supercomputer Center), funded by Ghent University, FWO and the Flemish Government – department EWI.

## Funding

F.B. was supported by Research Foundation – Flanders (FWO).

## Competing interests

The authors have no relevant financial or non-financial interests to disclose.

## Author contributions

All authors contributed to the study conception and design. Data collection and laboratory work were performed by FB. Data analyses were done by FB and FD. The first draft of the manuscript was written by FB and all authors commented on previous versions of the manuscript. All authors read and approved the final manuscript.

## Data availability

Scripts and data are made available on github (https://github.com/FemkeBatsleer/LandGenBembix) for review and will be published on zenodo upon acceptance.

## Supplementary information

S1 Landscape types_1.pdf: detailed description of the how landscape categories from original sources were combined and reclassified.

S2 Single category optimisation runs_2.pdf: results for the optimisation and bootstrap analyses of the 7 single category rasters for 4 independent runs.

S3 bootstrapsummary_westcoast_3.xlsx: full results of bootstrapping of all possible multi-categorical models after optimisation of the complete study area.

S4 dune area clusters_4.pdf: detailed results for the analyses repeated on the 4 dune area clusters situated within the complete study area.

S5 bootstrapsummary_areasdunes_5.xlsx: full results of the bootstrapping of all possible multi-categorical models after optimisation dune area clusters situated within the study area.

## Notes

https://github.com/FemkeBatsleer/LandGenBembix

### Competing Interest Statement

The authors have declared no competing interest.

### Summary of Updates

revisions based on peer-review comments; restructuring and revising text; updating figures; no analyses redone

