## Supplementary material for "Strong gene flow across an urbanised coastal landscape in a dune specialist digger wasp": S1 Landscape types_1.docx

We combined 2 types of landscape rasters, and a manually drawn polygon for the beach, resulting in a map of 7 landscape categories (Figure S1.3). For landscape categories within dune areas, we used detailed vegetation maps (Figure S1.1; resolution 0.4m) available for the region derived from remote sensing data with a classification machine learning technique (Bonte et al. 2021). For landscape categories outside dune areas, we used the ground cover map 2018 (Figure S1.2) provided by the Flemish government (BBK 1m resolution, source: [www.geopunt.be](http://www.geopunt.be)). Table S1.1 gives an overview how the source maps were reclassified. Figure S1.1, S1.2 and S1.3 give maps representing the two original sources and the resulting reclassified map.

Table S1.1: Original source landscape categories and their reclassified landscape types (and map color from Figure 1 in main manuscript and Figure S1.3).

| **Source** | **Source landscape category** | **Reclassified landscape category** | **Map Color** |
| --- | --- | --- | --- |
| Dune vegetation classification maps | Beachrose | Scrubs |  |
|  | Bucktorn | Scrubs |  |
|  | Nushgrass | Scrubs |  |
|  | Fixed | Trees |  |
|  | Grass | Open dune |  |
|  | Marram | Open dune |  |
|  | Marram/herb | Open dune |  |
|  | Moss | Open dune |  |
|  | Scrub/saltmarsh | Scrubs |  |
|  | Sand | Scrubs |  |
| Beach polygon | Beach | Beach |  |
| BBK | Buildings | Urbanised |  |
|  | Roads | Urbanised |  |
|  | Other asphalted | Urbanised |  |
|  | Railways | Urbanised |  |
|  | Water | Water |  |
|  | Other non-asphalted | Urbanised |  |
|  | Farmland | Agriculture |  |
|  | Grass, scrubs | Scrubs |  |
|  | Trees | Trees |  |
|  | Grass, scrubs (agricultural land) | Agriculture |  |
|  | Grass, scrubs (overlapping roads) | Urbanised |  |
|  | Trees (overlapping roads) | Urbanised |  |
|  | Grass, scrub (overlapping water) | Water |  |
|  | Trees (overlapping water) | Water |  |

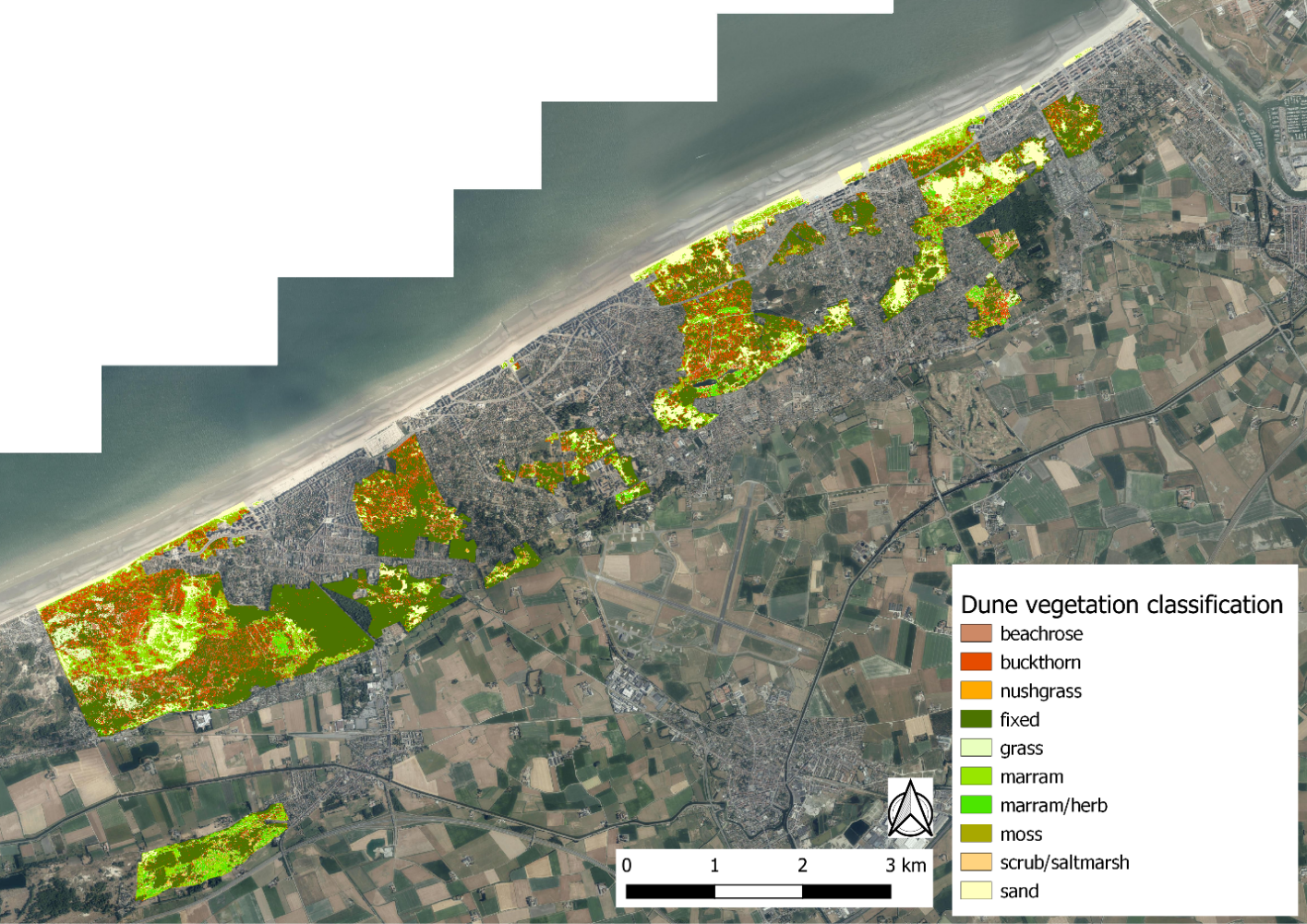
Figure S1.1: Dune vegetation classification map (see Bonte et al. 2021).
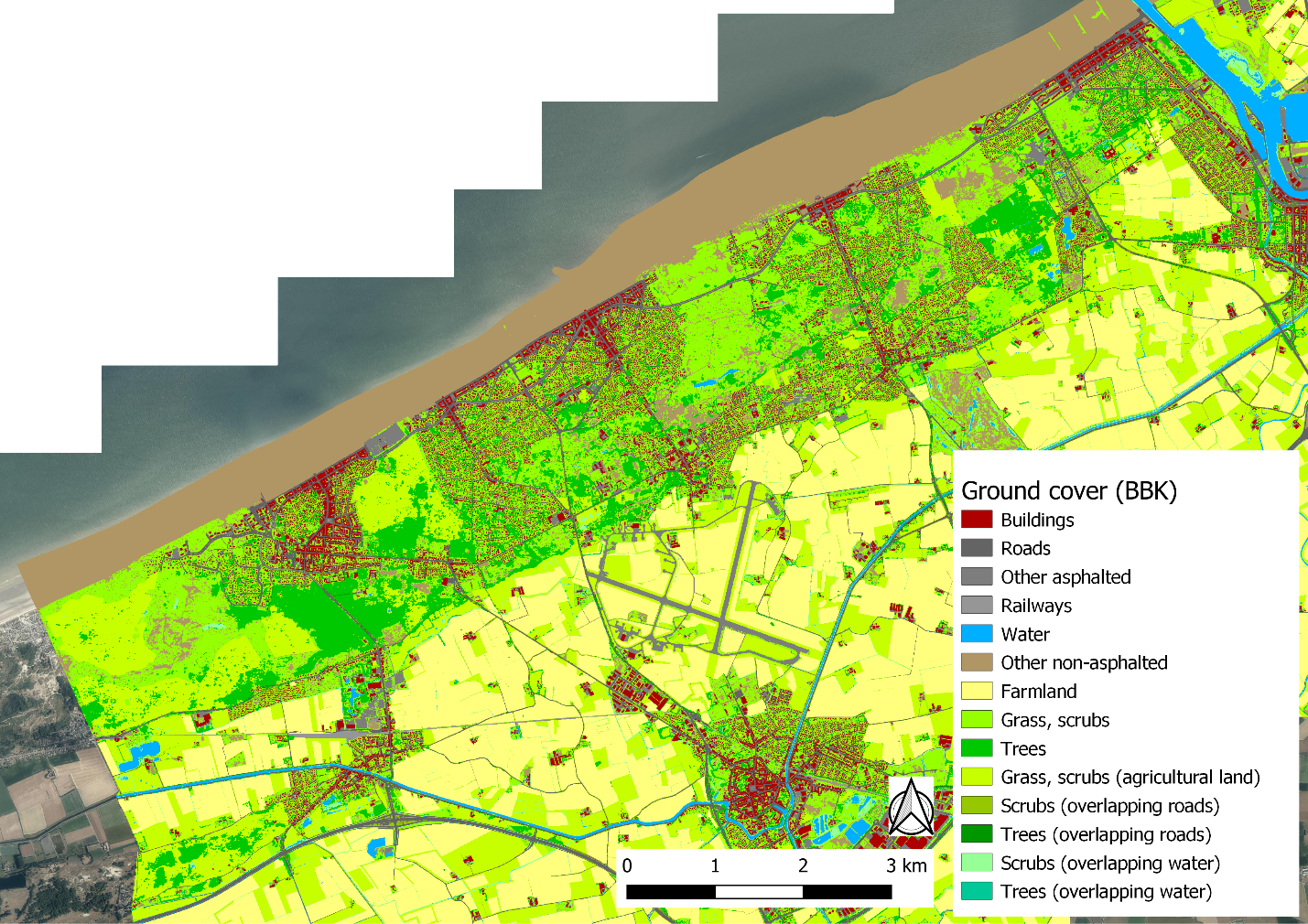
Figure S1.2: Ground cover map (BBK, source: [www.geopunt.be](http://www.geopunt.be)).

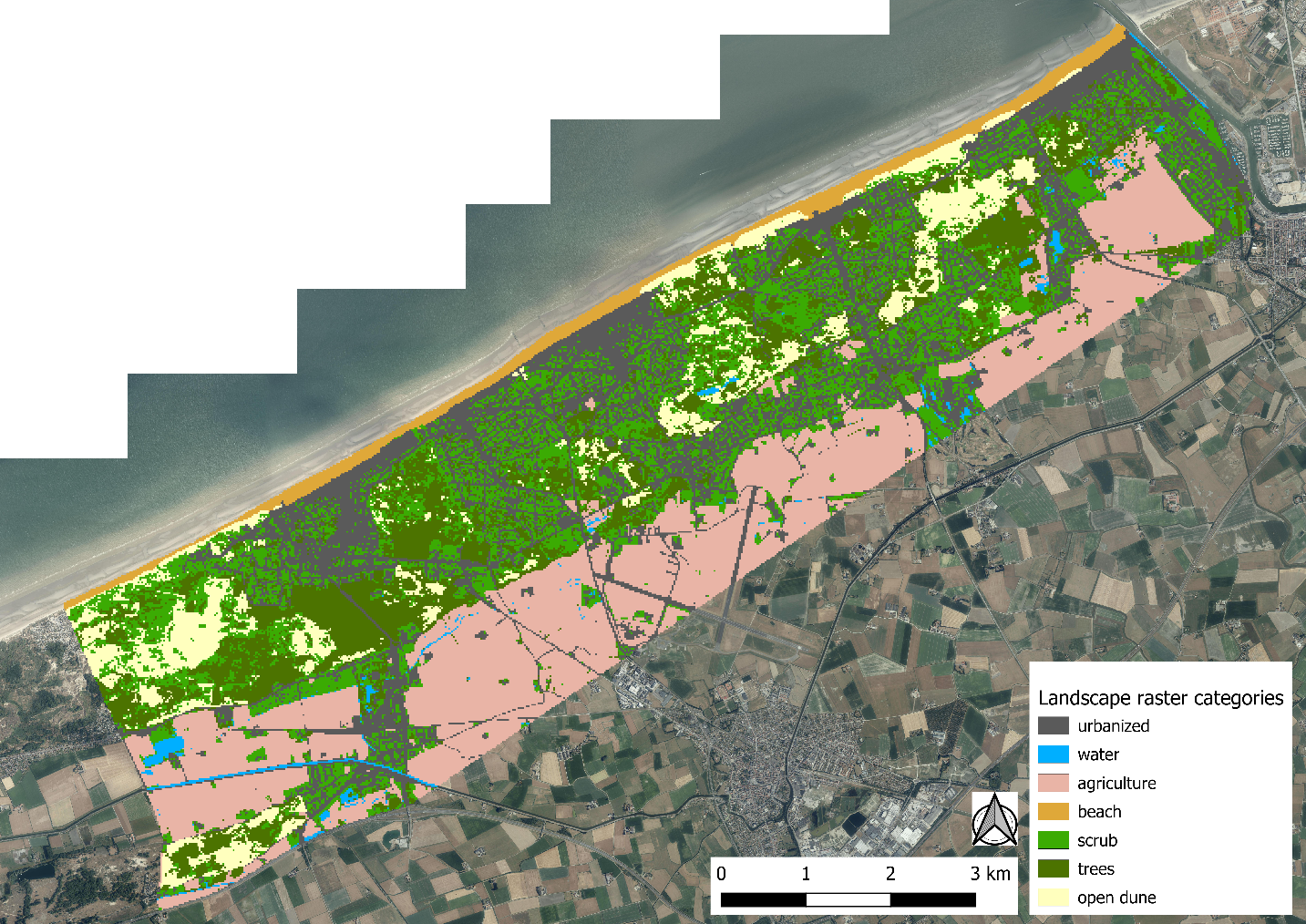
Figure S1.3: Reclassified landscape map.
