## Supplementary material for "Strong gene flow across an urbanised coastal landscape in a dune specialist digger wasp": S2 Single category optimisation runs_2.docx

Results of the optimisations of single category models after bootstrapping for four independent optimisation runs can be found in table S2.1-2.4 for the complete study.

For the complete study area, the four independent optimisation runs confirmed each other and the overall pattern. The *urban* model is the best supported model with a low resistance value (facilitating gene flow). Isolation-by-distance (*Distance* in table S2.1-S2.4) is the second best model with a ΔAICc < 2. Although, *beach* and *scrub* models were not stable and swapped between runs to either impeding or facilitating gene flow (table S2.1-S2.4). Results regarding those two categories should be considered inconclusive.

Table S2.1: Bootstrap results for the single category model optimisations for the complete study area, run 1.

| **Model** | **k** | **avg.AICc** | **ΔAICc** | **avg.weight** | **avg.R²m** | **avg.LL** | **n.top** | **Res.value. predictor** | **Res.value. other** |
| --- | --- | --- | --- | --- | --- | --- | --- | --- | --- |
| urban | 3 | 239681.26 | 0.00 | 0.39 | 0.00011 | -119837.60 | 610 | 1 | 2500 |
| Distance | 2 | 239682.55 | 1.29 | 0.20 | 0.00001 | -119839.26 | 339 | - | - |
| beach | 3 | 239684.58 | 3.32 | 0.07 | 0.00001 | -119839.26 | 0 | 2500 | 1 |
| water | 3 | 239684.61 | 3.35 | 0.07 | 0.00002 | -119839.27 | 0 | 2500 | 1 |
| trees | 3 | 239684.83 | 3.58 | 0.06 | 0.00001 | -119839.39 | 1 | 1 | 2500 |
| agric | 3 | 239684.93 | 3.68 | 0.06 | 0.00007 | -119839.44 | 0 | 2500 | 1 |
| scrub | 3 | 239684.94 | 3.69 | 0.07 | 0.00011 | -119839.44 | 25 | 2500 | 1 |
| opend | 3 | 239684.98 | 3.72 | 0.07 | 0.00010 | -119839.46 | 25 | 2500 | 1 |

Notes: 7 landscape categories compared in bootstrap analysis after optimisation of single category models. Urbanised (urban), beach, water, trees, agriculture (agric), scrub, open dune (opend). Distance is the isolation-by-distance null model: increasing genetic distance with increasing Euclidean geographic distance. Model, landscape category or univariate model; k, number of parameters; avg.AICc, average AICc across all bootstrap iterations; ΔAICc, difference in avg.AICc compared to the lowest avg.AICc (the best supported model); avg.weight, average Akaike weight across iterations; avg.R²m, average marginal R² across iterations; avg.LL, average log-likelihood across iterations; n.top, number of times the model was the top model across iterations; Res.value.predictor, the optimised resistance value for the focal landscape category; Res.value.other, optimised resistance value for all else (combined into one landscape variable). Models which have ΔAICc > 2 are coloured light grey.

Table S2.2: Bootstrap results for the single category model optimisations for the complete study area, run 2.

| **Model** | **k** | **avg.AICc** | **ΔAICc** | **avg.weight** | **avg.R²m** | **avg.LL** | **n.top** | **Res.value. predictor** | **Res.value. other** |
| --- | --- | --- | --- | --- | --- | --- | --- | --- | --- |
| urban | 3 | 239692.42 | 0.00 | 0.40 | 0.00011 | -119843.18 | 614 | 2500 | 1 |
| Distance | 2 | 239693.71 | 1.29 | 0.20 | 0.00001 | -119844.84 | 338 | NA | NA |
| beach | 3 | 239695.74 | 3.32 | 0.07 | 0.00001 | -119844.84 | 0 | 1 | 2500 |
| water | 3 | 239695.77 | 3.35 | 0.07 | 0.00002 | -119844.85 | 0 | 1 | 2500 |
| trees | 3 | 239696.00 | 3.59 | 0.06 | 0.00001 | -119844.97 | 2 | 2500 | 1 |
| agric | 3 | 239696.09 | 3.68 | 0.06 | 0.00007 | -119845.02 | 1 | 1 | 2500 |
| scrub | 3 | 239696.15 | 3.73 | 0.07 | 0.00010 | -119845.04 | 24 | 1 | 2500 |
| opend | 3 | 239696.22 | 3.81 | 0.07 | 0.00010 | -119845.08 | 21 | 1 | 2500 |

Notes: see table S2.1.

Table S2.3: Bootstrap results for the single category model optimisations for the complete study area, run 3.

| **Model** | **k** | **avg.AICc** | **ΔAICc** | **avg.weight** | **avg.R²m** | **avg.LL** | **n.top** | **Res.value. predictor** | **Res.value. other** |
| --- | --- | --- | --- | --- | --- | --- | --- | --- | --- |
| urban | 3 | 239699.10 | 0.00 | 0.39 | 0.00011 | -119846.52 | 616 | 2500 | 1 |
| Distance | 2 | 239700.40 | 1.29 | 0.19 | 0.00001 | -119848.18 | 333 | NA | NA |
| scrub | 3 | 239701.89 | 2.79 | 0.10 | 0.00002 | -119847.92 | 36 | 2500 | 1 |
| beach | 3 | 239702.43 | 3.32 | 0.07 | 0.00001 | -119848.18 | 0 | 1 | 2500 |
| water | 3 | 239702.46 | 3.36 | 0.07 | 0.00002 | -119848.20 | 0 | 1 | 2500 |
| trees | 3 | 239702.69 | 3.58 | 0.06 | 0.00001 | -119848.31 | 1 | 2500 | 1 |
| agric | 3 | 239702.78 | 3.68 | 0.06 | 0.00008 | -119848.36 | 0 | 1 | 2500 |
| opend | 3 | 239703.03 | 3.93 | 0.06 | 0.00009 | -119848.48 | 14 | 1 | 2500 |

Notes: see table S2.1.

Table S2.4: Bootstrap results for the single category model optimisations for the complete study area, run 4.

| **Model** | **k** | **avg.AICc** | **ΔAICc** | **avg.weight** | **avg.R²m** | **avg.LL** | **n.top** | **Res.value. predictor** | **Res.value. other** |
| --- | --- | --- | --- | --- | --- | --- | --- | --- | --- |
| urban | 3 | 239627.75 | 0.00 | 0.32 | 0.00011 | -119810.84 | 446 | 2500 | 1 |
| beach | 3 | 239628.03 | 0.28 | 0.28 | 0.00005 | -119810.98 | 340 | 2500 | 1 |
| Distance | 2 | 239629.18 | 1.43 | 0.15 | 0.00001 | -119812.57 | 176 | NA | NA |
| water | 3 | 239631.24 | 3.49 | 0.05 | 0.00002 | -119812.59 | 0 | 1 | 2500 |
| trees | 3 | 239631.44 | 3.69 | 0.05 | 0.00001 | -119812.69 | 2 | 2500 | 1 |
| opend | 3 | 239631.59 | 3.85 | 0.05 | 0.00010 | -119812.77 | 19 | 1 | 2500 |
| scrub | 3 | 239631.68 | 3.93 | 0.05 | 0.00009 | -119812.81 | 17 | 1 | 2500 |
| agric | 3 | 239631.56 | 3.81 | 0.04 | 0.00007 | -119812.75 | 0 | 1 | 2500 |

Notes: see table S2.1.
