## Supplementary material for "Strong gene flow across an urbanised coastal landscape in a dune specialist digger wasp": S4 dune area clusters_4.docx

For the four dune area clusters, results of the top 5 best supported models with optimisations of multi-categorical models (all possible combinations of landscape categories) after bootstrapping can be found in table S4.1 (full table in supplementary S5, including independent optimisation run 2). Figures similar to figure 2 are figures S4.2-S4.5 and table similar to table 2 is table S4.2 (full tables in supplementary S5).

Isolation-by-distance (IBD) is the overall best supported model for the dune area clusters, but there are a few exceptions and competing models (which have ΔAICc < 2). For Cabour, *agriculture* (impeding gene flow contrasting to the remaining landscape categories; table S4.1, Fig. S4.3) is the second best supported model, competing with IBD. For Westhoek, the combined model contrasting *beach.scrub.opend* to the remaining landscape categories is competing with IBD. These types of natural landscape (*beach*, *scrub*, *open dune*) are being barriers to gene flow (table S4.1, Fig. S4.2). For Doornpanne, the best supported model is *trees* contrasted to the remaining categories (facilitating gene flow; table S4.1, Fig. S4.4) with IBD a highly competing model. The four independent optimisation runs of single categories for the dune area clusters also confirm each other and the overall pattern of the multi-categorical optimisation runs (tables S4.3-S4.6). Several of the models with ΔAICc > 2 were not consistent in being either impeding or facilitating to gene flow across independent runs (table S4.3-S4.6). Thus, results regarding other categories then the best supported ones should be considered inconclusive on the scale of the four dune area clusters.

Table S4.1: Top 5 bootstrap results for the multi-categorical model optimisations for each dune area clusters.

| **Dune area** | **Model** | **k** | **avg.AICc** | **ΔAICc** | **avg.weight** | **avg.R²m** | **avg.LL** | **n.top** | **Res.value 1^st^** | **Res.value 2nd** | **Res.value 3rd** | **Res.value 4^th^** | **Res.value other** |
| --- | --- | --- | --- | --- | --- | --- | --- | --- | --- | --- | --- | --- | --- |
| Westhoek | Distance | 2 | 29848.91 | 0.00 | 0.18 | 0.00024 | -14922.41 | 514 | - | - | - | - | - |
|  | beach.scrub.opend | 5 | 29850.55 | 1.64 | 0.09 | 0.00184 | -14920.05 | 152 | 1338 | 2500 | 659 | - | 1 |
|  | scrub.opend | 4 | 29850.78 | 1.87 | 0.07 | 0.00140 | -14921.24 | 90 | 2500 | 451 | - | - | 1 |
|  | beach | 3 | 29851.00 | 2.09 | 0.06 | 0.00024 | -14922.41 | 0 | 2500 | - | - | - | 1 |
|  | urban.beach.scrub.opend | 6 | 29851.12 | 2.20 | 0.07 | 0.00163 | -14919.24 | 95 | 70 | 207 | 2500 | 526 | 1 |
| Cabour | Distance | 2 | 1496.54 | 0.00 | 0.19 | 0.00050 | -746.05 | 634 | - | - | - | - | - |
|  | agric | 3 | 1497.50 | 0.96 | 0.13 | 0.02864 | -745.29 | 289 | 2500 | - | - | - | 1 |
|  | opend | 3 | 1498.90 | 2.36 | 0.06 | 0.00184 | -745.99 | 19 | 2500 | - | - | - | 1 |
|  | trees | 3 | 1498.96 | 2.43 | 0.06 | 0.00086 | -746.02 | 14 | 2500 | - | - | - | 1 |
|  | water | 3 | 1499.02 | 2.48 | 0.05 | 0.00049 | -746.05 | 0 | 2500 | - | - | - | 1 |
| Doornpanne | trees | 3 | 9930.95 | 0.00 | 0.10 | 0.00089 | -4962.31 | 474 | 1 | - | - | - | 2500 |
|  | Distance | 2 | 9931.05 | 0.09 | 0.09 | 0.00019 | -4963.44 | 399 | - | - | - | - | - |
|  | opend | 3 | 9932.91 | 1.96 | 0.04 | 0.00119 | -4963.29 | 100 | 2500 | - | - | - | 1 |
|  | trees.opend | 4 | 9933.13 | 2.18 | 0.04 | 0.00098 | -4962.29 | 0 | 1 | 2500 | - | - | 1708 |
|  | beach.fixed | 4 | 9933.18 | 2.22 | 0.03 | 0.00089 | -4962.31 | 0 | 946 | 1 | - | - | 2500 |
| Teryde | Distance | 2 | 16509.20 | 0.00 | 0.13 | 0.00010 | -8252.54 | 944 | - | - | - | - | - |
|  | agric | 3 | 16511.32 | 2.12 | 0.04 | 0.00010 | -8252.54 | 0 | 2500 | - | - | - | 1 |
|  | beach | 3 | 16511.32 | 2.12 | 0.04 | 0.00010 | -8252.54 | 0 | 2500 | - | - | - | 1 |
|  | water | 3 | 16511.32 | 2.12 | 0.04 | 0.00010 | -8252.54 | 0 | 2500 | - | - | - | 1 |
|  | opend | 3 | 16511.68 | 2.47 | 0.04 | 0.00011 | -8252.72 | 1 | 1 | - | - | - | 1 |

Notes: 7 landscape categories compared in bootstrap analysis after optimisation of single category models. Urbanised (urban), beach, water, trees, agriculture (agric), scrub, open dune (opend). Distance is the isolation-by-distance null model: increasing genetic distance with increasing Euclidean geographic distance. Dune area, name of dune area cluster (Fig. 4.1); Predictor, landscape category or univariate model; k, number of parameters; avg.AICc, average AICc across all bootstrap iterations; ΔAICc, difference in avg.AICc compared to the lowest avg.AICc (the best supported model); avg.weight, average weight across iterations; avg.rank, average rank across iterations; avg.R²m, average marginal R² across iterations; avg.LL, average log-likelihood across iterations; n.top, number of times the model was the top model across iterations (does not sum op to 1000, as many more models were considered, see supplementary 5); Res.value 1^st^/2^nd^/3^rd^/4^th^, the optimised resistance value for the first/second/third/fourth mentioned landscape category (e.g. urban/beach.scrub/opend in the fifth row); Res.value other, optimised resistance value for all else (combined into one landscape variable).

Table S4.2: Summed Akaike weights per category for multivariate surfaces (multiple categories optimisations) for the dune area clusters.

| **Dune area** | **Category** | **Sum weights** | **Sum weights run 2** |
| --- | --- | --- | --- |
| Westhoek | scrub | 0.551 | 0.537 |
|  | opend | 0.465 | 0.460 |
|  | beach | 0.434 | 0.408 |
|  | trees | 0.283 | 0.264 |
|  | urban | 0.280 | 0.266 |
| Cabour | agric | 0.341 | 0.331 |
|  | water | 0.231 | 0.228 |
|  | scrub | 0.207 | 0.201 |
|  | opend | 0.188 | 0.191 |
|  | trees | 0.187 | 0.191 |
|  | urban | 0.187 | 0.187 |
| Doornpanne | trees | 0.408 | 0.417 |
|  | opend | 0.307 | 0.311 |
|  | water | 0.271 | 0.264 |
|  | beach | 0.244 | 0.235 |
|  | scrub | 0.228 | 0.233 |
|  | agric | 0.226 | 0.237 |
|  | urban | 0.222 | 0.222 |
| Ter Yde | scrub | 0.266 | 0.271 |
|  | agric | 0.264 | 0.261 |
|  | trees | 0.258 | 0.258 |
|  | water | 0.257 | 0.254 |
|  | beach | 0.257 | 0.255 |
|  | urban | 0.252 | 0.257 |
|  | opend | 0.249 | 0.248 |

Notes: the Akaike weight for a model (optimised surface) was added to the summation (Sum weights) if the focal category (Category) was in it.

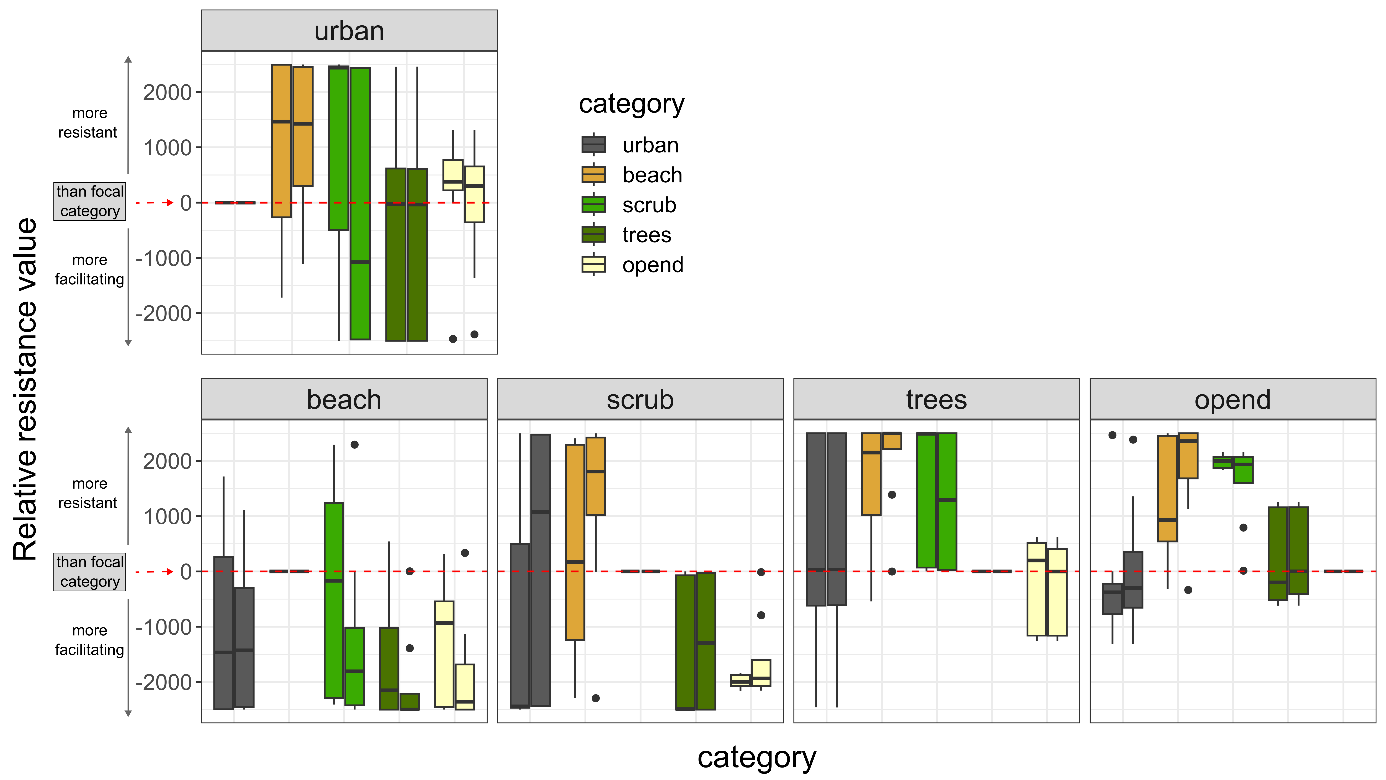

Figure S4.2: Relative resistance value from the optimisation of multivariate surfaces for dune area cluster Westhoek; similar to figure 2; The two independent runs are combined in as paired boxplots for each category in each panel (left run 1, right run 2).

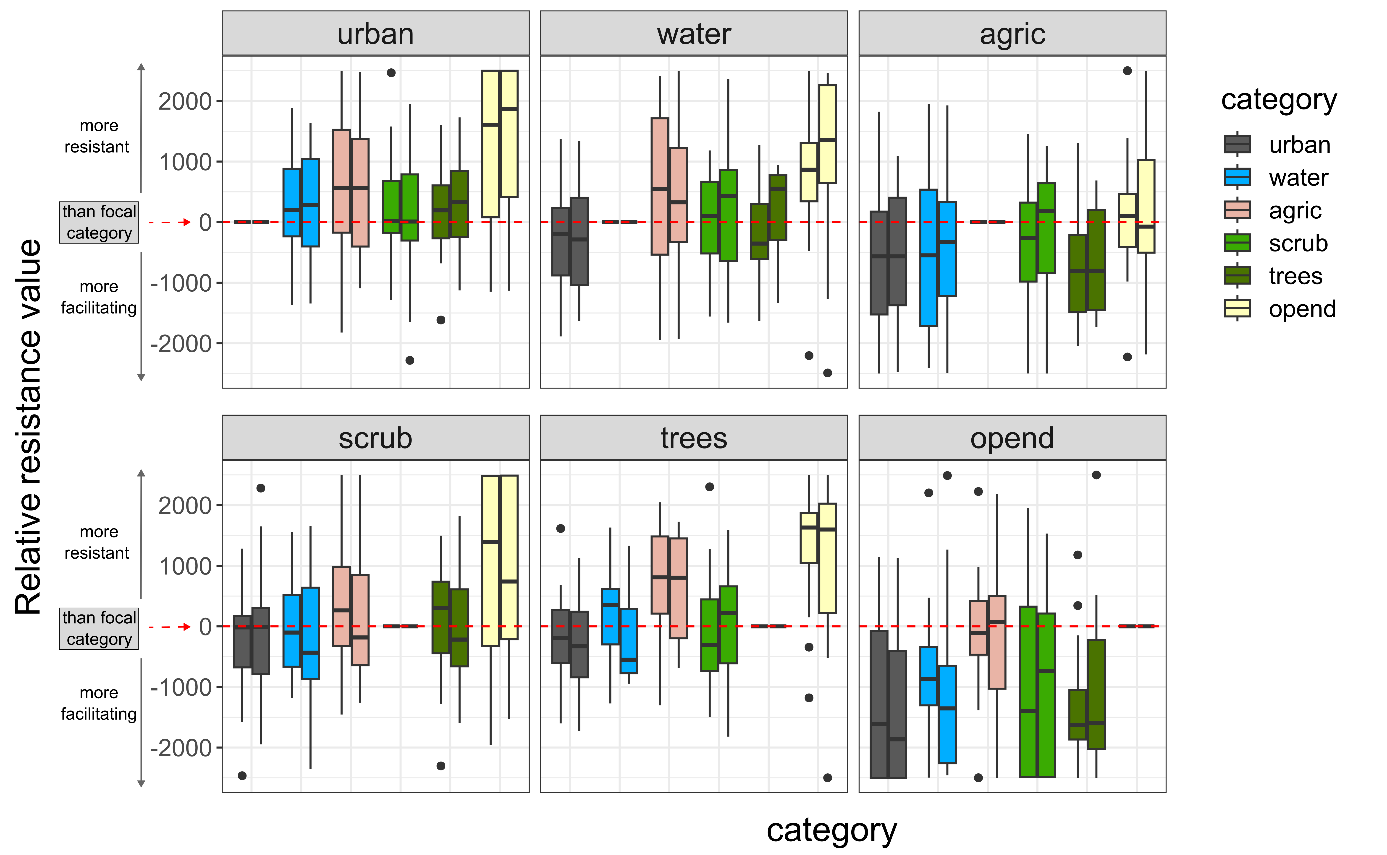

Figure S4.3: Relative resistance value from the optimisation of multivariate surfaces for dune area cluster Cabour; similar to figure 2; The two independent runs are combined in as paired boxplots for each category in each panel (left run 1, right run 2).

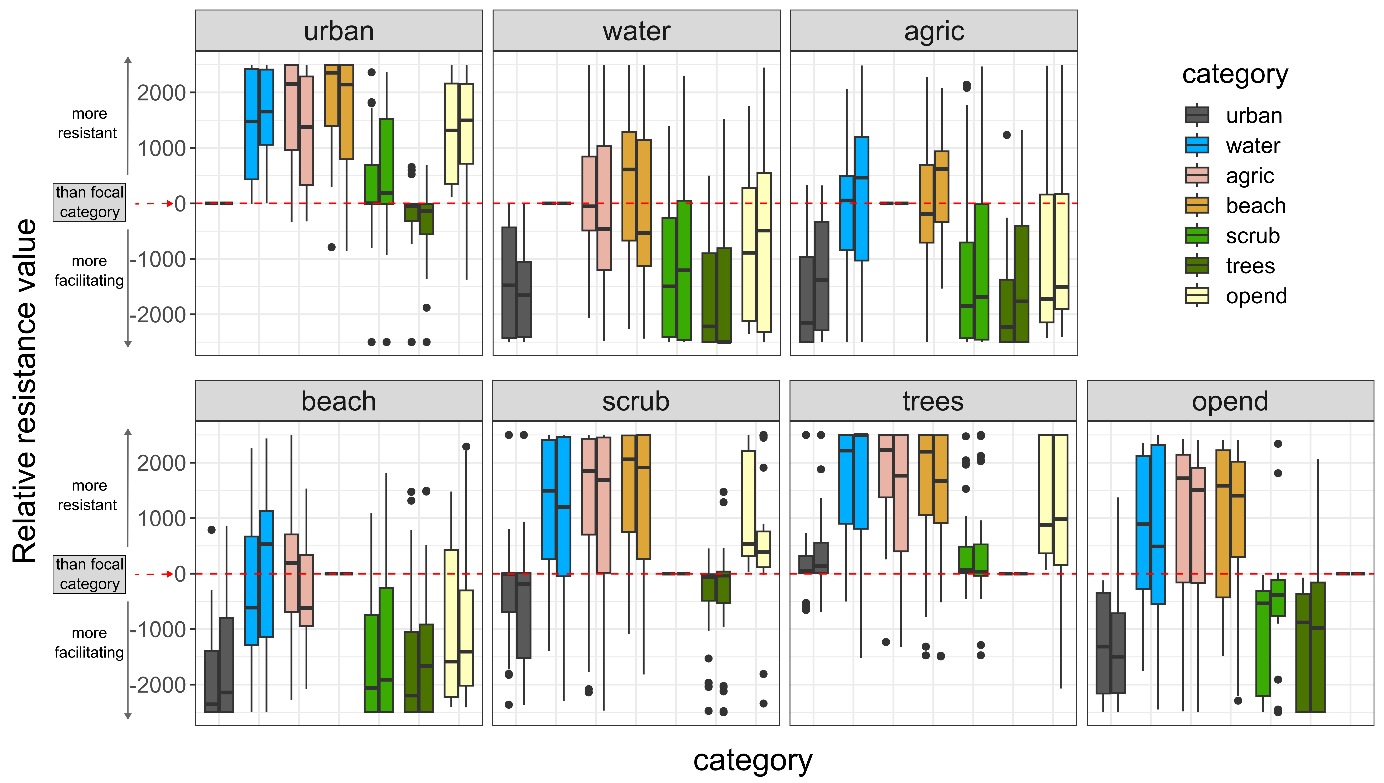

Figure S4.4: Relative resistance value from the optimisation of multivariate surfaces for dune area cluster Doornpanne; The two independent runs are combined in as paired boxplots for each category in each panel (left run 1, right run 2).

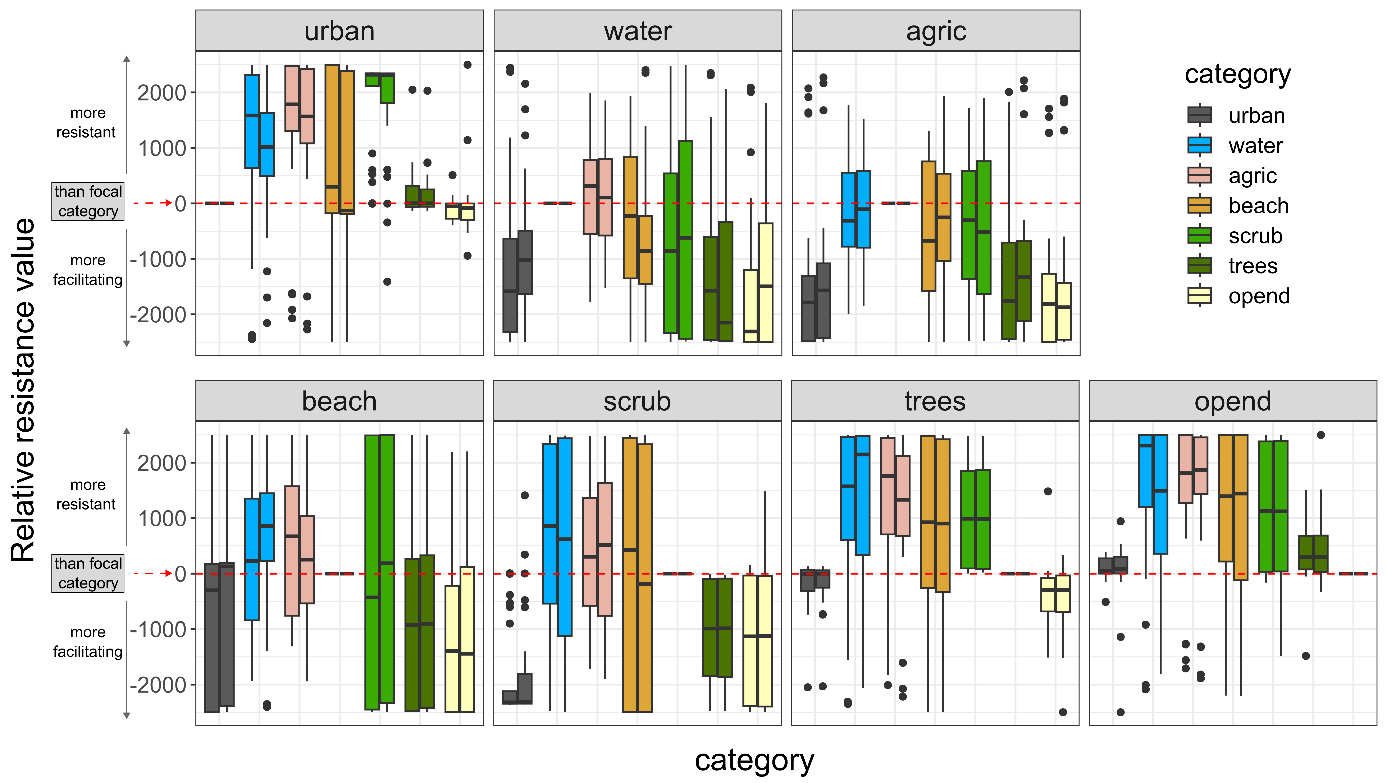
Figure S4.5: Relative resistance value from the optimisation of multivariate surfaces for dune area cluster Ter Yde; The two independent runs are combined in as paired boxplots for each category in each panel (left run 1, right run 2).

Table S4.3: Bootstrap results for the single category model optimisations for the dune area clusters, run 1.

| **Dune area** | **Model** | **k** | **avg.AICc** | **ΔAICc** | **avg.weight** | **avg.R2m** | **avg.LL** | **n.top** | **Res.value. predictor** | **Res.value. other** |
| --- | --- | --- | --- | --- | --- | --- | --- | --- | --- | --- |
| Westhoek | Distance | 2 | 29842.52 | 0.00 | 0.50 | 0.00025 | -14919.21 | 901 | NA | NA |
|  | beach | 3 | 29844.61 | 2.09 | 0.17 | 0.00025 | -14919.21 | 0 | 2500 | 1 |
|  | urban | 3 | 29845.76 | 3.25 | 0.11 | 0.00025 | -14919.79 | 15 | 1 | 2500 |
|  | scrub | 3 | 29847.22 | 4.70 | 0.07 | 0.00596 | -14920.52 | 5 | 2500 | 1 |
|  | trees | 3 | 29847.75 | 5.24 | 0.07 | 0.00074 | -14920.79 | 35 | 2500 | 1 |
|  | opend | 3 | 29847.98 | 5.47 | 0.07 | 0.00014 | -14920.90 | 44 | 1 | 2500 |
| Cabour | Distance | 2 | 1496.39 | 0.00 | 0.31 | 0.00049 | -745.97 | 661 | NA | NA |
|  | agric | 3 | 1497.41 | 1.02 | 0.22 | 0.02727 | -745.24 | 312 | 2500 | 1 |
|  | opend | 3 | 1498.77 | 2.38 | 0.10 | 0.00175 | -745.92 | 15 | 2500 | 1 |
|  | trees | 3 | 1498.83 | 2.44 | 0.10 | 0.00085 | -745.95 | 12 | 2500 | 1 |
|  | water | 3 | 1498.87 | 2.48 | 0.09 | 0.00049 | -745.97 | 0 | 2500 | 1 |
|  | urban | 3 | 1498.87 | 2.48 | 0.09 | 0.00049 | -745.97 | 0 | 2500 | 1 |
|  | scrub | 3 | 1498.87 | 2.48 | 0.09 | 0.00049 | -745.97 | 0 | 2500 | 1 |
| Doornpanne | trees | 3 | 9930.23 | 0.00 | 0.27 | 0.00089 | -4961.95 | 447 | 1 | 2500 |
|  | Distance | 2 | 9930.37 | 0.13 | 0.23 | 0.00018 | -4963.10 | 415 | NA | NA |
|  | opend | 3 | 9931.99 | 1.76 | 0.13 | 0.00135 | -4962.83 | 126 | 2500 | 1 |
|  | agric | 3 | 9932.53 | 2.30 | 0.08 | 0.00018 | -4963.10 | 0 | 2500 | 1 |
|  | beach | 3 | 9932.53 | 2.30 | 0.08 | 0.00018 | -4963.10 | 0 | 2500 | 1 |
|  | water | 3 | 9932.54 | 2.31 | 0.08 | 0.00017 | -4963.11 | 0 | 2500 | 1 |
|  | urban | 3 | 9933.01 | 2.78 | 0.07 | 0.00015 | -4963.35 | 12 | 2500 | 1 |
|  | scrub | 3 | 9933.11 | 2.87 | 0.06 | 0.00013 | -4963.39 | 0 | 1 | 2500 |
| Teryde | Distance | 2 | 16499.84 | 0.00 | 0.31 | 0.00010 | -8247.86 | 953 | NA | NA |
|  | water | 3 | 16501.96 | 2.12 | 0.11 | 0.00010 | -8247.86 | 0 | 2500 | 1 |
|  | beach | 3 | 16501.96 | 2.12 | 0.11 | 0.00010 | -8247.86 | 0 | 2500 | 1 |
|  | agric | 3 | 16501.96 | 2.12 | 0.11 | 0.00010 | -8247.86 | 0 | 2500 | 1 |
|  | scrub | 3 | 16502.34 | 2.51 | 0.10 | 0.00056 | -8248.05 | 20 | 2500 | 1 |
|  | opend | 3 | 16502.35 | 2.51 | 0.09 | 0.00011 | -8248.05 | 3 | 1 | 2500 |
|  | trees | 3 | 16502.39 | 2.55 | 0.10 | 0.00050 | -8248.07 | 20 | 2500 | 1 |
|  | urban | 3 | 16502.50 | 2.67 | 0.09 | 0.00009 | -8248.13 | 4 | 1 | 2500 |

Notes: 7 landscape categories compared in bootstrap analysis after optimisation of single category models. Urbanised (urban), beach, water, trees, agriculture (agric), scrub, open dune (opend). Distance is the isolation-by-distance null model: increasing genetic distance with increasing Euclidean geographic distance. Dune area, name of dune area cluster (Fig. 1); Predictor, landscape category or univariate model; k, number of parameters; avg.AICc, average AICc across all bootstrap iterations; ΔAICc, difference in avg.AICc compared to the lowest avg.AICc (the best supported model); avg.weight, average weight across iterations; avg.R²m, average marginal R² across iterations; avg.LL, average log-likelihood across iterations; n.top, number of times the model was the top model across iterations; Res.value.predictor, the optimised resistance value for the focal landscape category; Res.value.other, optimised resistance value for all else (combined into one landscape variable). Models which have ΔAICc > 2 are coloured light grey.

Table S4.4: Bootstrap results for the single category model optimisations for the dune area clusters, run 2.

| **Dune area** | **Model** | **k** | **avg.AICc** | **ΔAICc** | **avg.weight** | **avg.R2m** | **avg.LL** | **n.top** | **Res.value. predictor** | **Res.value. other** |
| --- | --- | --- | --- | --- | --- | --- | --- | --- | --- | --- |
| Westhoek | Distance | 2 | 29846.00 | 0.00 | 0.48 | 0.00024 | -14920.96 | 894 | NA | NA |
|  | beach | 3 | 29848.09 | 2.09 | 0.17 | 0.00024 | -14920.96 | 0 | 1 | 2500 |
|  | urban | 3 | 29849.22 | 3.22 | 0.11 | 0.00024 | -14921.52 | 10 | 2500 | 1 |
|  | trees | 3 | 29850.00 | 3.99 | 0.10 | 0.00032 | -14921.91 | 37 | 2500 | 1 |
|  | scrub | 3 | 29850.48 | 4.47 | 0.07 | 0.00634 | -14922.15 | 13 | 1 | 2500 |
|  | opend | 3 | 29851.26 | 5.25 | 0.08 | 0.00014 | -14922.54 | 46 | 2500 | 1 |
| Cabour | Distance | 2 | 1496.64 | 0.00 | 0.31 | 0.00051 | -746.10 | 647 | NA | NA |
|  | agric | 3 | 1497.68 | 1.03 | 0.21 | 0.02757 | -745.38 | 311 | 1 | 2500 |
|  | opend | 3 | 1498.97 | 2.33 | 0.10 | 0.00196 | -746.02 | 28 | 1 | 2500 |
|  | trees | 3 | 1499.06 | 2.42 | 0.10 | 0.00091 | -746.07 | 14 | 1 | 2500 |
|  | water | 3 | 1499.12 | 2.48 | 0.09 | 0.00051 | -746.10 | 0 | 1 | 2500 |
|  | scrub | 3 | 1499.13 | 2.48 | 0.09 | 0.00051 | -746.10 | 0 | 1 | 2500 |
|  | urban | 3 | 1499.12 | 2.48 | 0.09 | 0.00051 | -746.10 | 0 | 1 | 2500 |
| Doornpanne | trees | 3 | 9932.44 | 0.00 | 0.28 | 0.00090 | -4963.06 | 470 | 2500 | 1 |
|  | Distance | 2 | 9932.55 | 0.11 | 0.23 | 0.00018 | -4964.19 | 424 | NA | NA |
|  | opend | 3 | 9934.44 | 2.00 | 0.12 | 0.00113 | -4964.06 | 94 | 1 | 2500 |
|  | agric | 3 | 9934.71 | 2.27 | 0.08 | 0.00018 | -4964.19 | 0 | 1 | 2500 |
|  | beach | 3 | 9934.71 | 2.27 | 0.08 | 0.00018 | -4964.19 | 0 | 1 | 2500 |
|  | water | 3 | 9934.72 | 2.29 | 0.08 | 0.00018 | -4964.20 | 0 | 1 | 2500 |
|  | urban | 3 | 9935.24 | 2.80 | 0.07 | 0.00014 | -4964.46 | 12 | 1 | 2500 |
|  | scrub | 3 | 9935.33 | 2.89 | 0.06 | 0.00013 | -4964.50 | 0 | 2500 | 1 |
| Teryde | Distance | 2 | 16505.96 | 0.00 | 0.30 | 0.00011 | -8250.92 | 945 | NA | NA |
|  | scrub | 3 | 16508.05 | 2.09 | 0.11 | 0.00009 | -8250.90 | 14 | 2500 | 1 |
|  | agric | 3 | 16508.08 | 2.12 | 0.10 | 0.00011 | -8250.92 | 0 | 1 | 2500 |
|  | beach | 3 | 16508.08 | 2.12 | 0.10 | 0.00011 | -8250.92 | 0 | 1 | 2500 |
|  | water | 3 | 16508.08 | 2.12 | 0.10 | 0.00011 | -8250.92 | 0 | 1 | 2500 |
|  | trees | 3 | 16508.50 | 2.54 | 0.10 | 0.00052 | -8251.13 | 27 | 1 | 2500 |
|  | opend | 3 | 16508.47 | 2.51 | 0.09 | 0.00012 | -8251.11 | 6 | 2500 | 1 |
|  | urban | 3 | 16508.60 | 2.64 | 0.09 | 0.00010 | -8251.18 | 8 | 2500 | 1 |

Notes: see table S4.3.

Table S4.5: Bootstrap results for the single category model optimisations for the dune area clusters, run 3.

| **Dune area** | **Model** | **k** | **avg.AICc** | **ΔAICc** | **avg.weight** | **avg.R2m** | **avg.LL** | **n.top** | **Res.value. predictor** | **Res.value. other** |
| --- | --- | --- | --- | --- | --- | --- | --- | --- | --- | --- |
| Westhoek | Distance | 2 | 29844.56 | 0.00 | 0.47 | 0.00024 | -14920.24 | 915 | NA | NA |
|  | beach | 3 | 29846.65 | 2.09 | 0.17 | 0.00024 | -14920.24 | 0 | 1 | 2500 |
|  | urban | 3 | 29846.98 | 2.41 | 0.14 | 0.00023 | -14920.40 | 0 | 1 | 2500 |
|  | scrub | 3 | 29849.17 | 4.60 | 0.07 | 0.00623 | -14921.49 | 13 | 1 | 2500 |
|  | opend | 3 | 29850.05 | 5.48 | 0.07 | 0.00013 | -14921.93 | 37 | 2500 | 1 |
|  | trees | 3 | 29849.76 | 5.20 | 0.07 | 0.00075 | -14921.79 | 35 | 1 | 2500 |
| Cabour | Distance | 2 | 1496.92 | 0.00 | 0.31 | 0.00051 | -746.24 | 641 | NA | NA |
|  | agric | 3 | 1497.90 | 0.98 | 0.22 | 0.02905 | -745.49 | 325 | 1 | 2500 |
|  | opend | 3 | 1499.33 | 2.41 | 0.10 | 0.00168 | -746.21 | 19 | 1 | 2500 |
|  | trees | 3 | 1499.36 | 2.44 | 0.10 | 0.00088 | -746.22 | 15 | 1 | 2500 |
|  | water | 3 | 1499.40 | 2.48 | 0.09 | 0.00051 | -746.24 | 0 | 1 | 2500 |
|  | scrub | 3 | 1499.40 | 2.48 | 0.09 | 0.00052 | -746.24 | 0 | 1 | 2500 |
|  | urban | 3 | 1499.40 | 2.48 | 0.09 | 0.00051 | -746.24 | 0 | 1 | 2500 |
| Doornpanne | trees | 3 | 9929.87 | 0.00 | 0.28 | 0.00089 | -4961.77 | 470 | 2500 | 1 |
|  | Distance | 2 | 9929.95 | 0.08 | 0.23 | 0.00019 | -4962.90 | 410 | NA | NA |
|  | opend | 3 | 9931.73 | 1.86 | 0.12 | 0.00123 | -4962.70 | 106 | 1 | 2500 |
|  | agric | 3 | 9932.12 | 2.25 | 0.08 | 0.00019 | -4962.90 | 0 | 1 | 2500 |
|  | beach | 3 | 9932.12 | 2.25 | 0.08 | 0.00019 | -4962.90 | 0 | 1 | 2500 |
|  | water | 3 | 9932.13 | 2.26 | 0.08 | 0.00018 | -4962.90 | 0 | 1 | 2500 |
|  | urban | 3 | 9932.64 | 2.77 | 0.07 | 0.00016 | -4963.16 | 14 | 1 | 2500 |
|  | scrub | 3 | 9933.06 | 3.19 | 0.06 | 0.00048 | -4963.37 | 0 | 1 | 2500 |
| Teryde | Distance | 2 | 16501.49 | 0.00 | 0.30 | 0.00010 | -8248.69 | 935 | NA | NA |
|  | agric | 3 | 16503.62 | 2.12 | 0.10 | 0.00010 | -8248.69 | 0 | 1 | 2500 |
|  | water | 3 | 16503.62 | 2.12 | 0.10 | 0.00010 | -8248.69 | 0 | 1 | 2500 |
|  | beach | 3 | 16503.62 | 2.12 | 0.10 | 0.00010 | -8248.69 | 0 | 1 | 2500 |
|  | trees | 3 | 16504.01 | 2.51 | 0.10 | 0.00052 | -8248.88 | 20 | 1 | 2500 |
|  | scrub | 3 | 16503.93 | 2.44 | 0.10 | 0.00064 | -8248.84 | 26 | 1 | 2500 |
|  | opend | 3 | 16503.97 | 2.47 | 0.09 | 0.00012 | -8248.86 | 9 | 2500 | 1 |
|  | urban | 3 | 16504.15 | 2.66 | 0.09 | 0.00009 | -8248.95 | 10 | 2500 | 1 |

Notes: see table S4.3.

Table S4.6: Bootstrap results for the single category model optimisations for the dune area clusters, run 4.

| **Dune area** | **Model** | **k** | **avg.AICc** | **ΔAICc** | **avg.weight** | **avg.R2m** | **avg.LL** | **n.top** | **Res.value. predictor** | **Res.value. other** |
| --- | --- | --- | --- | --- | --- | --- | --- | --- | --- | --- |
| Westhoek | Distance | 2 | 29850.13 | 0.00 | 0.46 | 0.00025 | -14923.02 | 914 | NA | NA |
|  | beach | 3 | 29852.22 | 2.09 | 0.16 | 0.00025 | -14923.02 | 0 | 1 | 2500 |
|  | urban | 3 | 29852.57 | 2.44 | 0.14 | 0.00024 | -14923.19 | 1 | 1 | 2500 |
|  | trees | 3 | 29854.25 | 4.12 | 0.09 | 0.00033 | -14924.03 | 34 | 2500 | 1 |
|  | scrub | 3 | 29854.87 | 4.74 | 0.07 | 0.00604 | -14924.34 | 6 | 1 | 2500 |
|  | opend | 3 | 29855.62 | 5.49 | 0.08 | 0.00014 | -14924.72 | 45 | 2500 | 1 |
| Cabour | Distance | 2 | 1496.18 | 0.00 | 0.31 | 0.00055 | -745.87 | 628 | NA | NA |
|  | agric | 3 | 1497.15 | 0.96 | 0.22 | 0.02881 | -745.11 | 328 | 1 | 2500 |
|  | opend | 3 | 1498.54 | 2.36 | 0.10 | 0.00199 | -745.81 | 27 | 1 | 2500 |
|  | trees | 3 | 1498.60 | 2.42 | 0.10 | 0.00095 | -745.84 | 17 | 1 | 2500 |
|  | water | 3 | 1498.66 | 2.48 | 0.09 | 0.00055 | -745.87 | 0 | 1 | 2500 |
|  | scrub | 3 | 1498.67 | 2.48 | 0.09 | 0.00055 | -745.87 | 0 | 1 | 2500 |
|  | urban | 3 | 1498.67 | 2.48 | 0.09 | 0.00054 | -745.87 | 0 | 1 | 2500 |
| Doornpanne | trees | 3 | 9929.32 | 0.00 | 0.29 | 0.00092 | -4961.50 | 480 | 2500 | 1 |
|  | Distance | 2 | 9929.52 | 0.20 | 0.23 | 0.00018 | -4962.68 | 403 | NA | NA |
|  | opend | 3 | 9931.30 | 1.99 | 0.12 | 0.00123 | -4962.49 | 107 | 1 | 2500 |
|  | agric | 3 | 9931.68 | 2.37 | 0.08 | 0.00018 | -4962.68 | 0 | 1 | 2500 |
|  | beach | 3 | 9931.68 | 2.37 | 0.08 | 0.00018 | -4962.68 | 0 | 1 | 2500 |
|  | water | 3 | 9931.70 | 2.38 | 0.08 | 0.00018 | -4962.69 | 0 | 1 | 2500 |
|  | urban | 3 | 9932.21 | 2.89 | 0.07 | 0.00015 | -4962.94 | 10 | 1 | 2500 |
|  | scrub | 3 | 9932.28 | 2.96 | 0.06 | 0.00013 | -4962.98 | 0 | 2500 | 1 |
| Teryde | Distance | 2 | 16508.31 | 0.00 | 0.30 | 0.00010 | -8252.09 | 943 | NA | NA |
|  | beach | 3 | 16510.43 | 2.12 | 0.11 | 0.00010 | -8252.09 | 0 | 1 | 2500 |
|  | water | 3 | 16510.43 | 2.12 | 0.11 | 0.00010 | -8252.09 | 0 | 1 | 2500 |
|  | agric | 3 | 16510.43 | 2.12 | 0.11 | 0.00010 | -8252.09 | 0 | 1 | 2500 |
|  | trees | 3 | 16510.85 | 2.55 | 0.10 | 0.00049 | -8252.30 | 16 | 1 | 2500 |
|  | scrub | 3 | 16510.81 | 2.51 | 0.10 | 0.00057 | -8252.28 | 27 | 1 | 2500 |
|  | opend | 3 | 16510.78 | 2.48 | 0.09 | 0.00012 | -8252.27 | 5 | 2500 | 1 |
|  | urban | 3 | 16510.95 | 2.65 | 0.09 | 0.00010 | -8252.35 | 9 | 2500 | 1 |

Notes: see table S4.3.
